# Widely used GWAS methods can be poorly suited to SNP-level localization under diffuse polygenic architecture in livestock

**DOI:** 10.64898/2026.07.20.739621

**Authors:** Xuesong Wang, Junjian Wang, Francesco Tiezzi, Yijian Huang, Wen Huang, Christian Maltecca, Jicai Jiang

## Abstract

In livestock populations, genome-wide association studies (GWAS) can produce strong, apparently localized associations even when no truly discrete nearby causal effect exists. This occurs because small effective population sizes, strong family structure, long-range linkage disequilibrium (LD), and diffuse polygenic architecture can cause the effects of many variants to accumulate and be captured jointly across broad genomic intervals, making variant-level associations difficult to interpret biologically. Using real pig genotypes, we constructed a benchmark in which phenotypes were simulated under diffuse polygenic architecture across a genome partitioned into alternating effect and null windows, with central-null regions (at least 1 Mb away from effect-containing regions) positioned to detect long-range LD-driven signal propagation. We evaluated nine configurations of six GWAS methods (BOLT-LMM, REGENIE, fastGWA, FarmCPU, BLINK, and SLEMM) under this architecture. The central finding is that strong associations, of the kind normally read as evidence of nearby moderate- or large-effect variants, are produced by many of these methods even though the simulated signal is distributed across many tiny effects and cannot be localized to any single variant. The methods differed sharply in the extent of locus-level spillover: several produced large numbers of genome-wide significant loci within central-null regions, whereas the full-GRM mixed-model benchmark (SLEMM) produced no genome-wide significant loci in central-null regions. These results show that, under a highly polygenic architecture with livestock-like LD, GWAS tool choice has major consequences for biological interpretation. When the goal is to localize biologically meaningful signals rather than to flag association peaks that may merely reflect tiny effects accumulated through LD across a broad block, methods that control long-range LD spillover should be prioritized.

## Introduction

Livestock genetic breeding plays a vital role in agriculture by improving productivity, profitability, and resilience to environmental change, and advances in genomics have enabled the widespread use of genome-wide association studies (GWAS) in cattle, sheep, pigs, and poultry^1–16^. GWAS is applied not only to detect genotype–phenotype associations, but also to localize trait-associated genomic regions for downstream biological interpretation, including candidate gene prioritization, fine-mapping, colocalization with transcriptomic data, pathway analysis, and functional validation^17–26^. The biological value of a GWAS signal therefore depends not only on its statistical significance, but critically on whether it reliably reflects a nearby, localized genetic effect (an assumption that underlies most standard post-GWAS analyses)^27,28^.

However, this assumption is systematically challenged by two features of livestock GWAS that interact to undermine signal interpretability. First, livestock populations have small effective population sizes, strong family structure, and extensive long-range LD arising from their shared demographic history, which preserves long identical-by-descent segments and maintains LD over broad genomic distances, producing highly related populations with large half-sib and full-sib families^29–34^. Empirical studies in cattle and pig have confirmed that variant-level associations frequently reflect broad linked haplotypic segments rather than narrowly localized causal effects, and that long-range LD can render many correlated but noncausal variants apparently associated^35–45^. Second, most economically relevant traits in livestock are complex quantitative traits governed by highly polygenic architectures, in which phenotypic variation is collectively attributable to large numbers of variants each with individually small effect sizes^26,46^. Under the polygenic and omnigenic frameworks^26,46^, the per-variant effects of such traits are expected to fall below conventional genome-wide significance thresholds and should not, in principle, be detectable as discrete significant signals in standard GWAS^47,48^. Indeed, sub-significant variants have been shown to account for a substantial proportion of the heritability of complex traits^26^, confirming that genome-wide significant signals alone represent only a partial and potentially misleading view of the underlying genetic architecture.

These two features interact in a particularly problematic way: long-range LD in livestock genomes can propagate and amplify diffuse polygenic effects across broadly correlated genomic regions, inflating association statistics at linked markers and producing signals that achieve genome-wide significance not because they tag a discrete nearby causal variant, but because long-range LD allows them to aggregate effects from many causal variants^41,43,46,49^. Consequently, many GWAS methods may transform what is fundamentally a diffuse, small-effect polygenic signal into apparently localized association peaks that mimic genuine discrete genetic effects. Such signals are prone to misinterpretation in downstream analyses, because they achieve statistical significance through LD-mediated accumulation of polygenic background rather than through proximity to a causal variant^50–53^. A recent theoretical framework formalizes this mechanism, showing that under the full-GRM mixed model the per-SNP non-centrality parameter is bounded by an effective marker-count ceiling M_e_ = 4N_e_L set by the effective population size and genome length (in Morgans), and that leave-one-chromosome-out (LOCO) and full-GRM detect fundamentally different signal types: LOCO captures LD-aggregated block-level signals while full-GRM detects only SNP-level excess effects above the polygenic baseline^54^.

This concern is method-dependent, because GWAS approaches differ fundamentally in how they partition phenotypic variation into tested marker effects and background polygenic covariance^55–59^. Linear mixed models absorb background variation through a random genetic component; whole-genome regression methods estimate all marker effects jointly with shrinkage; and multi-locus methods iteratively reassign markers between fixed and background components. Under diffuse polygenic architectures, these differences in model structure determine how readily small effects carried on correlated haplotypes are captured indirectly through LD, producing marginal associations that may reflect aggregate linked variation rather than a single, discrete causal signal^41,60^. Judging GWAS methods solely by the number of detected associations therefore risks overstating biological interpretability if sharper-looking peaks arise from LD-tagged polygenic background rather than genuine local resolution^41^.

We therefore asked not which method detects the greatest number of associations, but which methods preserve interpretable localization and which instead transform diffuse polygenic background into apparently localized but misleading association peaks. To address this question, we used pigs as an empirical example while focusing on the broader problem of GWAS interpretability in livestock populations. Using real pig genotypes to represent livestock-like genomic structure, we designed a benchmark in which phenotypes were simulated under a diffuse polygenic architecture across predefined signal and null genomic regions, allowing us to quantify spillover beyond true signal boundaries and compare a diverse set of GWAS methods that differ in how they model background signal. These included mixed-model, whole-genome regression, and multi-locus approaches such as BOLT-LMM^61^, REGENIE^62^, fastGWA^56^, FarmCPU^59^, BLINK^63^, and SLEMM^64^. Our main result is that, under diffuse polygenic effects in livestock-like genomes, several widely used methods can produce strong association signals in central-null regions that are misleading for localization, whereas the dense whole-genome mixed-model SLEMM produced no genome-wide significant loci in central-null regions.

## Materials and Methods

### Genomic data and simulation design

A total of 31,301 pigs genotyped with the PorcineSNP60 BeadChip (Illumina Inc., San Diego, CA, USA) and provided by Smithfield Premium Genetics (Roanoke Rapids, NC, USA) served as the genomic resource for this study. Sequence-level imputation was performed with SWIM 1.0, and quality control retained variants with MAF ≥ 1%, Hardy–Weinberg equilibrium *P* ≥ 1 × 10⁻⁹, and INFO score ≥ 0.8, yielding approximately 11.7 million variants. To reduce computational burden, sequence variants were thinned by retaining markers at least 20 kb apart on each chromosome using PLINK2 (--bp-space 20000). The resulting set was then merged with all directly genotyped chip markers, producing a final panel of approximately 135,358 SNPs for all subsequent analyses. The retained marker panel exhibited extensive long-range linkage disequilibrium extending over megabase scales (Figure 1).

**Figure 1.**
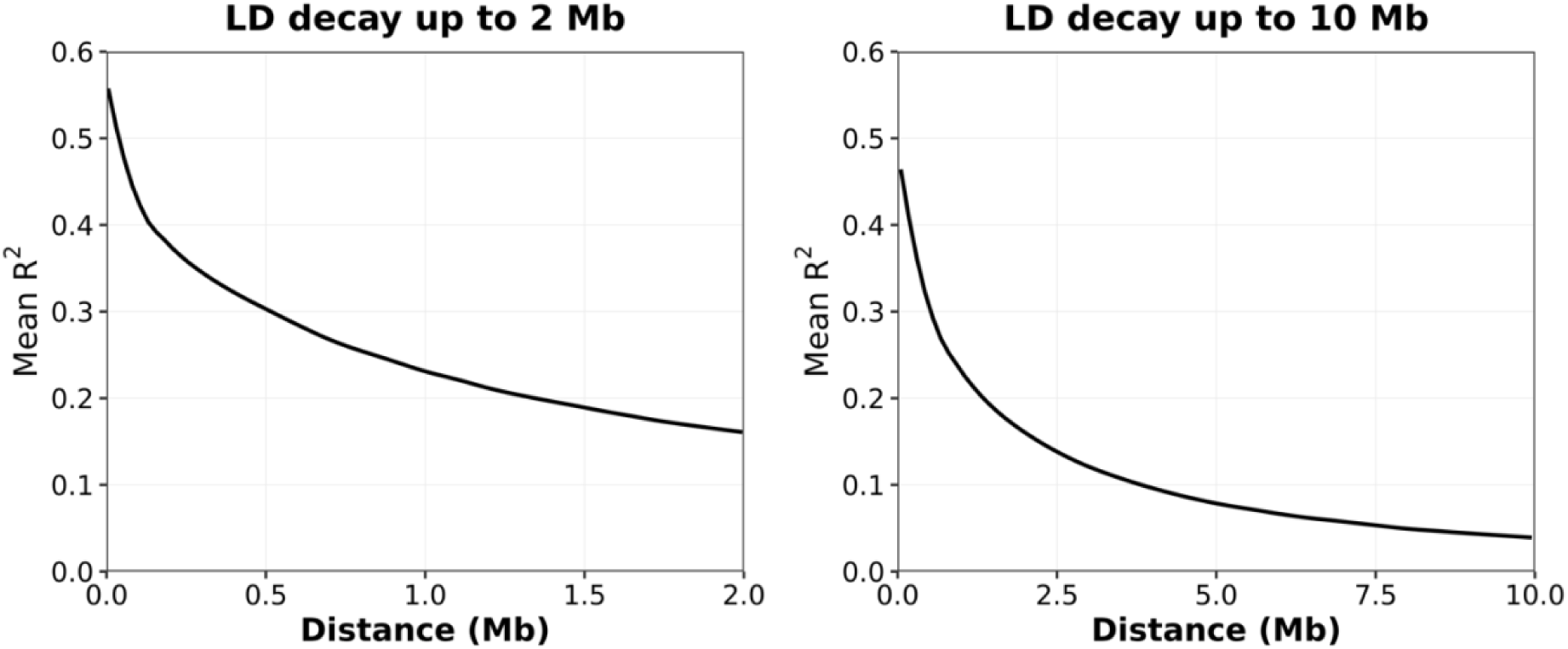
Long-range linkage disequilibrium extends over megabase scales in the pig population. Mean pairwise linkage disequilibrium (LD), measured as the squared Pearson correlation coefficient (*r*²) between SNP genotypes, computed with PLINK. The left panel shows LD decay up to 2 Mb, and the right panel extends the analysis to 10 Mb. Although LD decays with distance, appreciable correlation persists over several megabases, demonstrating the extensive long-range LD characteristic of this pig population.

Ten independent phenotype replicates were simulated under a polygenic additive model with total narrow-sense heritability fixed at *h*^2^ = 0.5 Polygenic effects were assigned to 10,000 SNPs randomly sampled from the polygenic-effect windows, with effect sizes independently drawn from 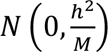, thereby restricting nonzero genetic effects to these regions rather than distributing them across the genome. Importantly, the locations of these nonzero-effect SNPs were fixed across all 10 phenotype replicates, whereas the SNP effect sizes and residual errors were independently redrawn from the same distributions in each replicate. A subset of 20,000 individuals was retained for all subsequent GWAS analyses, and 10 independently simulated phenotypes were used to assess the consistency of method performance across replicates.

Because individual effect sizes were randomly sampled, realized per-SNP contributions varied across replicates. Across the 10 simulated phenotypes, the largest realized per-SNP proportion of variance explained (PVE) ranged from 0.063% to 0.093% (12.5–18.5 times the nominal mean of 0.005%), corresponding to marginal non-centrality parameters of 12.5–18.5 and only 2.8%– 12.5% power at the genome-wide significance threshold. The expected number of causal variants detectable through their own marginal effects follows directly from the simulation design. The non-centrality parameter of a causal variant’s single-marker test is *δ* = *Nα*^2^ with 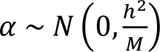, where *α* is the additive effect size of the causal variant. Consequently, *δ* is itself a random variable rather than a fixed quantity. With *N* = 20,000, ℎ^2^ = 0.5, and *M* = 10,000, the scaling factor 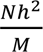 equals 1 exactly, so *δ* follows a chi-square distribution with one degree of freedom and has mean 1. Conditional on *δ*, the test statistic follows the non-central chi-square distribution 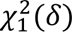. Averaging over the distribution of 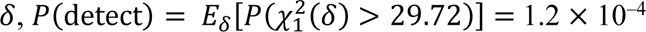 where 29.72 is the critical value corresponding to the genome-wide significance threshold (*P* = 5 × 10⁻^8^) Thus, although each simulated phenotype was controlled by 10,000 causal variants, only approximately 1.2 per replicate were expected to reach genome-wide significance through their own effect.

To evaluate the spatial propagation of association signals, chromosomes were partitioned into alternating polygenic-effect and null windows, with each 2 Mb polygenic-effect window followed by a 4 Mb null window, forming a recurring 6 Mb pattern along each chromosome. Within each null window, a central 2 Mb region (referred to as the central-null region throughout) was further defined, positioned 1 Mb away from the boundaries of adjacent causal windows. This design ensures that central-null regions are spatially separated from causal regions and minimally influenced by local LD at window boundaries, providing a framework for evaluating long-range LD-induced spillover across GWAS methods. To quantify long-range propagation of association signals into effect-free regions, we calculated a locus-level spillover metric using the central 2 Mb null regions. Each central-null region was partitioned into consecutive 500 kb windows, consistent with the locus definition used elsewhere in this study. A 500 kb window was considered a spillover locus if it contained at least one SNP reaching genome-wide significance (*P* ≤ 5 × 10⁻⁸). For each GWAS method and phenotype replicate, the spillover-locus rate was calculated as the number of significant central-null 500 kb windows divided by the total number of central-null 500 kb windows. Such loci represent apparent association signals generated by long-range LD with causal variants in the effect windows rather than by any local causal variant. The chromosomal window structure is illustrated in Figure 2. Additionally, for fine-scale evaluation, each null region was further partitioned into 50 kb windows and indexed by their distance from the nearest causal boundary (1, 2, …, 40 from each side). This indexing scheme enables systematic assessment of spillover-locus rates as a function of genomic distance from causal regions. For each 50 kb window, we calculated the observed-to-expected ratio of significant SNPs as the number of observed significant SNPs divided by the expected number under the nominal significance threshold (the number of SNPs tested within the window multiplied by α = 5 × 10⁻⁸). Ratios were subsequently averaged across windows with the same distance index to characterize the spatial extent of LD-driven signal propagation.

**Figure 2.**
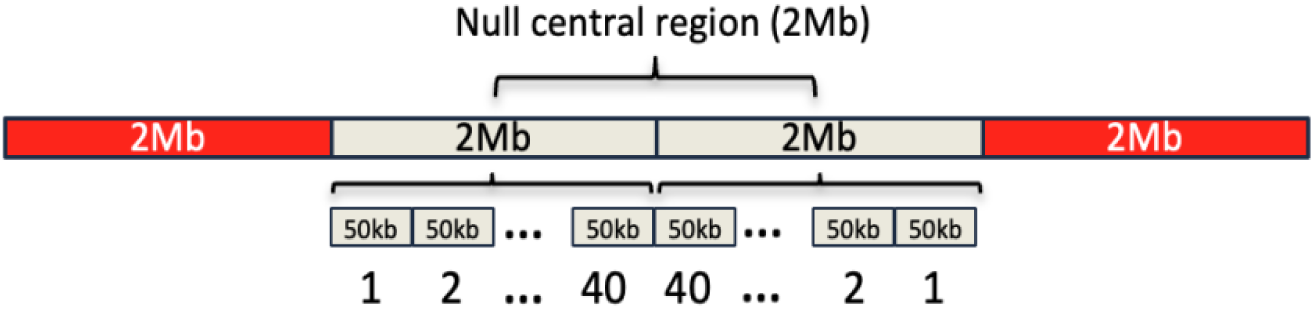
Genomic window design for analysis. The genome was divided into repeating 2 Mb polygenic-effect windows and 4 Mb null windows. Red regions indicate polygenic-effect windows containing the 10,000 nonzero-effect SNPs used to simulate phenotypes (ℎ^2^ = 0.5), and gray regions indicate null windows with no simulated effects. Within each null window, a central 2 Mb region (the central-null region) was defined to ensure separation from adjacent effect windows. Each null window was then partitioned symmetrically into 50 kb bins and indexed by distance from the nearest polygenic-effect boundary (1, 2, …, 40 from each side), allowing spillover-locus rates to be evaluated as a function of distance from effect regions.

Phenotypes were simulated under a standard linear additive genetic model:

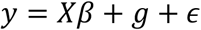

where:

- *y* ∈ ℝ*^n^* is the vector of phenotypes,
- *Xβ* represents fixed effects (intercept only in this study),
- 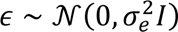 is the residual error,
- *g* = *Zα* is the vector of genetic effects.

where:

- *Z* ∈ ℝ*^n^*^×*m*^ is the standardized genotype matrix,
- α ∈ ℝ*^m^* is the vector of SNP effect sizes.

A total of 10,000 SNPs were randomly selected and assigned nonzero effects drawn from a normal distribution:

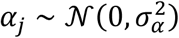

The remaining SNPs had zero effect. Effect sizes were scaled such that the total narrow-sense heritability was:

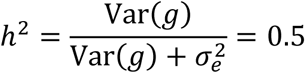

### GWAS methods evaluated

We benchmarked six GWAS methods (BOLT-LMM, REGENIE, fastGWA, FarmCPU, BLINK, and SLEMM), representing a range of commonly used association-testing frameworks. Ordinary linear regression without kinship correction was not included because uncorrected association testing is well known to produce severely inflated test statistics in livestock populations with extensive family structure and long-range linkage disequilibrium (LD), making it an inappropriate baseline for this setting^55,65–68^. We also evaluated alternative configurations for selected tools. REGENIE was run with both its default LOCO model and a LOCO-disabled model. For the GAPIT-based methods FarmCPU and BLINK^69^, both Kansas and NYC Manhattan plot outputs were assessed. Kansas reflects the standard multi-step framework and typically yields more discrete association peaks, whereas NYC applies additional iterative refinement and often produces broader, more continuous regional signals^70^. Thus, the benchmark comprised six core methods, with additional comparisons across alternative configurations for REGENIE, FarmCPU, and BLINK.

BOLT-LMM was run using BOLT-LMM v2.5 with the options --lmm, --LDscoresUseChip, and --numThreads 16. Analyses were conducted separately for each of the 10 simulated phenotypes using the PLINK dataset containing 135,358 SNPs and 31,301 genotyped individuals, of which 20,000 phenotyped individuals were retained for analysis. No additional covariates were included beyond the intercept term. No genetic map was provided, so physical positions were used. BOLT-LMM evaluated both infinitesimal and non-infinitesimal models internally, but because the non-infinitesimal model did not substantially improve fit, association results were taken from the infinitesimal mixed-model implementation.

REGENIE was run using REGENIE v4.1.2 in its standard two-step framework for each of the 10 simulated phenotypes, with --qt, --bsize 1000, and --threads 16. In the default LOCO implementation, step 1 fit whole-genome ridge regression models and generated chromosome-specific prediction files, and step 2 used the corresponding _pred.list files for single-marker testing. In the LOCO-disabled implementation, step 1 was run with --print-prs, and step 2 was run with --use-prs and the corresponding _prs.list files, so that genome-wide predictors included markers from the chromosome under test rather than excluding them. Thus, REGENIE was benchmarked under two distinct configurations: LOCO-enabled and LOCO-disabled.

fastGWA was run in GCTA v1.94.1 using --fastGWA-mlm. A full SNP-derived genomic relationship matrix (GRM) was first constructed from the PLINK genotype data with --make-grm and then converted to a sparse GRM with --make-bK-sparse 0.05. This procedure retained only pairwise relatedness estimates greater than 0.05, with smaller off-diagonal GRM elements set to zero, and the resulting sparse GRM was used for mixed-model association testing of each phenotype replicate.

FarmCPU and BLINK were implemented in GAPIT v3.5 using phenotype and genotype inputs derived from the same underlying PLINK dataset. For each method, both Kansas and NYC outputs were retained for evaluation and treated as alternative configurations of the same method rather than as separate GWAS methods. FarmCPU iteratively alternates between fixed- and random-effect models, using the random-effect step to jointly evaluate selected pseudo-QTNs and control confounding. BLINK, by contrast, uses an iterative fixed-effect framework without an explicit random-effect term, iteratively selecting associated markers and updating them as fixed covariates.

SLEMM v0.90.1 was used as the full-GRM mixed-model GWAS benchmark via its slemm_gwa routine. SLEMM implements the full-GRM mixed model and closely approximates the exact full-GRM results obtained with GCTA MLMA^55^ on these pig data^71^. For each of the 10 simulated phenotypes, a null linear mixed model was first fit with slemm --lmm using the phenotype file, the PLINK genotype dataset, and a SNP annotation file (16 threads). Chromosome-wise association testing was then performed with slemm_gwa for chromosomes 1– 18, and per-chromosome outputs were merged into a single genome-wide result file. The comparative analysis reported here uses the slemm_gwa results.

At the model level, BOLT-LMM and fastGWA are linear mixed-model approaches, with fastGWA using a sparse GRM for computational efficiency. REGENIE employs a two-step whole-genome regression framework. FarmCPU alternates between fixed- and random-effect modeling with selected pseudo-QTNs, whereas BLINK uses an iterative fixed-effect procedure without an explicit random effect. SLEMM is a full-GRM (dense, genome-wide) linear mixed model that uses window-based SNP weighting to model the polygenic background; unlike fastGWA it does not sparsify the GRM, and unlike BOLT-LMM and REGENIE it does not apply LOCO.

A uniform genome-wide significance threshold of 5 × 10⁻^8^ was applied across all methods.

### Theoretical NCP calculation

The theoretical non-centrality parameter (NCP) was computed for each of the 135,358 tested SNPs according to Equation 13 of Jiang^54^, using only the genotypes and the known simulated architecture, with no parameters fitted to the GWAS results.

Equation 13 expresses the LOCO non-centrality as the product of the LOCO coefficient, the sample size multiplied by the per-SNP causal effect variance, and the causal LD score.

The causal LD score of each tested SNP was computed as the sum of squared genotype correlations between that SNP and all 10,000 causal variants on the same chromosome, evaluated in the same 20,000 animals used for GWAS, retaining the self term when the tested SNP was itself causal. Correlations were computed blockwise from standardized genotypes and verified against PLINK --r2.

The per-SNP causal effect variance was *h*²/*m* = 0.5/10,000 = 5 × 10⁻⁵. Because *N* = 20,000, the product of the sample size and the per-SNP effect variance equals 1 exactly, so the predicted NCP reduces to the product of the LOCO coefficient and the causal LD score.

The LOCO coefficient (Equation 12 of Jiang^54^) was computed for each SNP from the leave-one-chromosome-out covariance matrix, which combined a genomic relationship matrix built from all 135,358 SNPs excluding the focal chromosome with a residual term. This background matches the one used by the LOCO methods. Chromosome-specific variance components were set to their expected values under the simulation design: each chromosome’s heritability was *h*² multiplied by its share of the 10,000 causal variants, and the leave-one-chromosome-out heritability was the remainder. Genotypes were centered and scaled by the square root of 2pq, matching the standardization used to simulate the phenotypes.

## Results

### Heritability estimation

To assess how well standard variance-component estimation recovers the simulated heritability under this architecture, we applied fastGWA-REML to estimate the SNP-based heritability 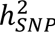 for each simulated phenotype replicate. Despite a simulation design heritability of 0.5, fastGWA-REML consistently returned substantially lower estimates across all replicates (mean = 0.145, range: 0.135–0.153), recovering only approximately 29% of the simulated heritability. In contrast, SLEMM’s dense-GRM GREML recovered essentially the full simulated heritability (mean ≈ 0.529, range: 0.506–0.550). This dense-versus-sparse contrast, consistent across replicates, identifies GRM sparsification, rather than the trait architecture or the estimator itself, as the direct cause of the downward bias. Because fastGWA approximates the full GRM by retaining only sparse kinship elements above a relatedness threshold, it captures only a fraction of the genome-wide polygenic signal, while the diffuse contribution of the many small-effect variants distributed across the genome and captured only through distant marker correlations is largely discarded. The missing heritability in this context is therefore not a biological phenomenon but a consequence of GRM sparsification under a highly polygenic architecture, and it implies that the polygenic background modeled during association testing by fastGWA is also incomplete, with direct consequences for the inflation and spillover examined below.

### Apparent association signals across GWAS methods

Comparison of GWAS results across methods revealed substantial differences in both the number and genomic distribution of significant associations, while preserving broadly similar phenotype-specific patterns (Figure 3). Across all 10 simulated phenotypes, the number of significant SNPs and the number of detected loci, defined as non-overlapping 500 kb windows containing at least one significant SNP, varied markedly by method. Methods that model genome-wide polygenic effects with dense, leave-one-chromosome-out predictors, particularly BOLT-LMM and REGENIE_LOCO, consistently detected far more significant SNPs and loci than the other approaches, often yielding thousands of loci per phenotype. By contrast, fastGWA detected fewer loci, whereas BLINK and FarmCPU, which rely on iterative multi-locus selection, were more restrictive and detected fewer of both. Notably, REGENIE_woLOCO detected substantially fewer loci than REGENIE_LOCO, showing that the contrast is methodological.

**Figure 3.**
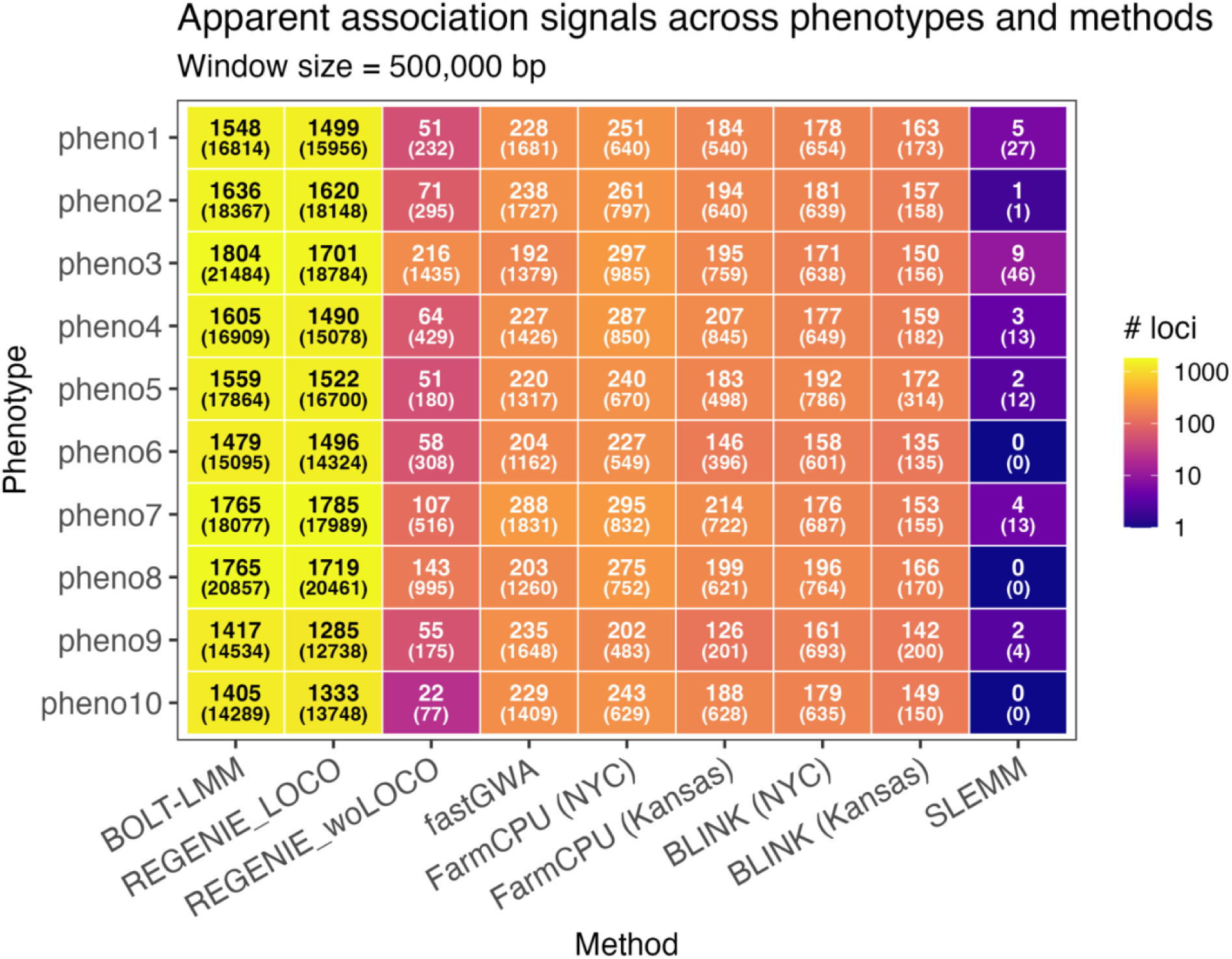
Genome-wide significant loci across the entire genome, including simulated effect and null regions, for 10 simulated phenotypes and nine method configurations. Detected loci were defined as non-overlapping 500 kb genomic windows containing at least one significant SNP. In each heatmap cell, the number at the top indicates the number of detected loci, and the number in parentheses at the bottom indicates the total number of significant SNPs. Colors represent the number of detected loci. REGENIE_LOCO and REGENIE_woLOCO denote REGENIE with and without LOCO, respectively.

The Manhattan plots (Figure 4) show that these differences are not only quantitative but also spatial. Complete Manhattan plots showing all tested SNPs for every method configuration separately for each simulated phenotype are provided in Supplementary Figures S1-S10. BOLT-LMM and REGENIE_LOCO produced extensive clusters of significant SNPs spanning broad genomic regions and forming visually prominent peaks that resemble localized association signals. The remaining methods generally produced fewer and more fragmented peaks, although they still generated discrete genomic clusters rather than a pattern that visibly reflected the underlying diffuse architecture. Thus, despite differences in stringency, several widely used GWAS methods transformed a genome-wide tiny-effect architecture into apparently localized variant-level signals.

**Figure 4.**
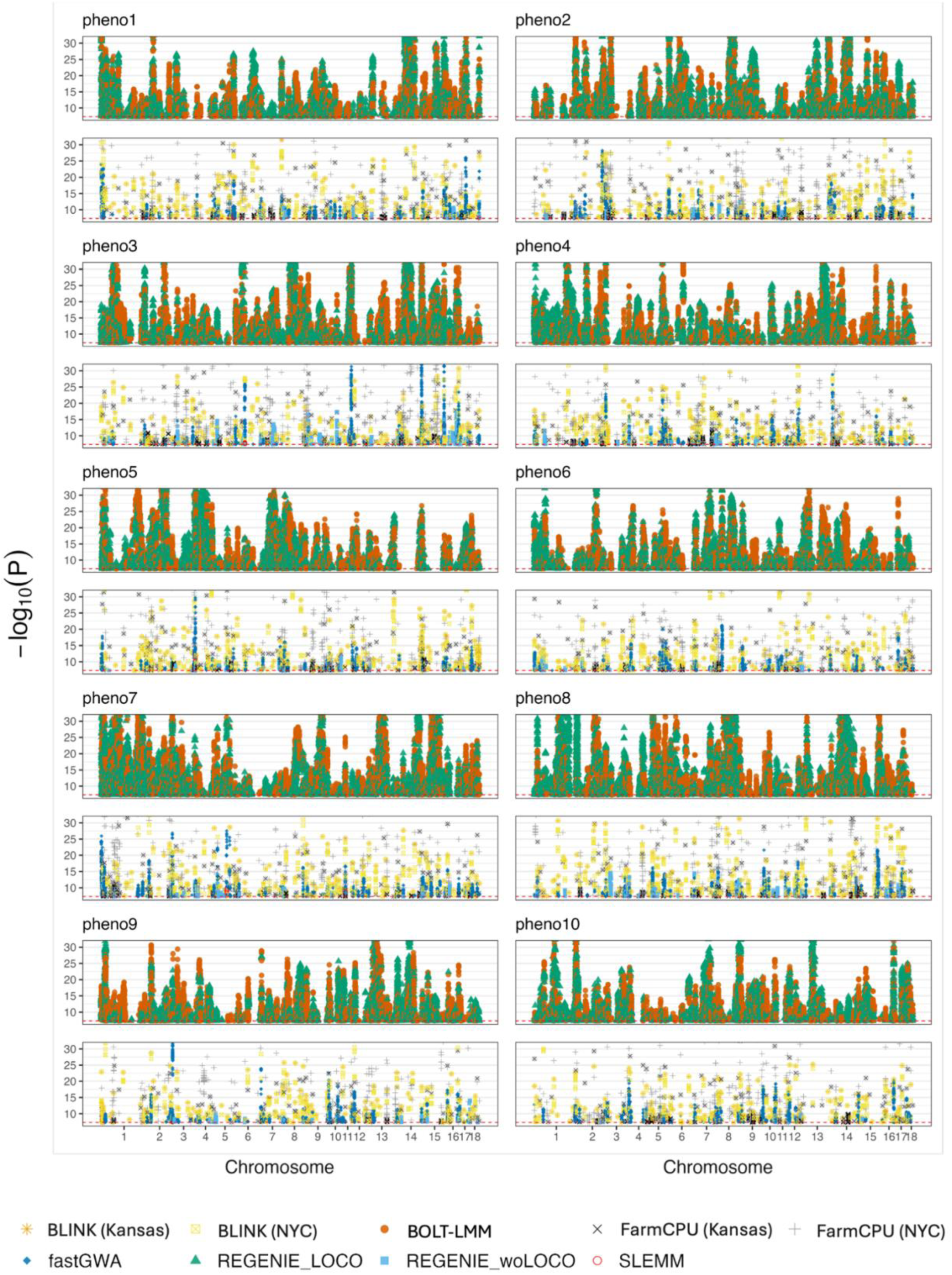
Manhattan plots of significant SNPs across 10 simulated phenotypes (shown in two stacked panels for clarity). The upper panel displays BOLT-LMM and REGENIE_LOCO, the lower panel shows the remaining methods.

Locus-level overlap analysis further showed that these apparent signals were not consistently localized to the same genomic regions across methods (Figure 5). The highest concordance was between configurations of the same method (BLINK (Kansas) versus BLINK (NYC) and FarmCPU (Kansas) versus FarmCPU (NYC)), and between distinct methods sharing a leave-one-chromosome-out strategy (BOLT-LMM versus REGENIE_LOCO). In contrast, overlap across broader methodological classes was generally modest, indicating that different methods often highlighted different sets of significant loci for the same phenotype. SLEMM showed near-zero overlap with the other methods, consistent with its highly conservative behavior and very limited number of detected loci.

**Figure 5.**
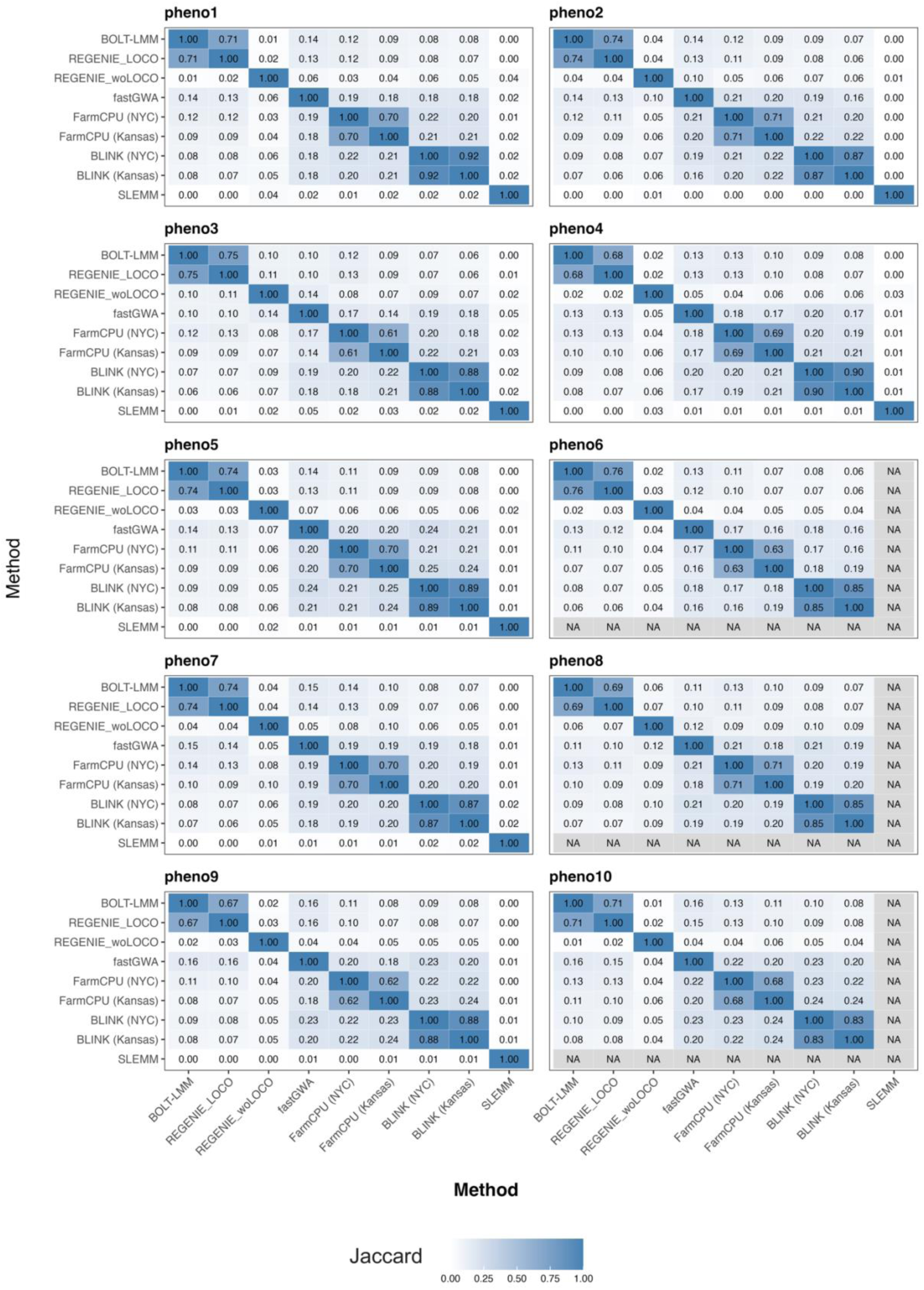
Locus-level overlap among GWAS methods across 10 simulated phenotypes (measured using the Jaccard index). Loci were defined as non-overlapping 500 kb genomic windows containing at least one significant SNP. Each panel corresponds to a phenotype, and each cell shows the pairwise Jaccard index between two configurations, quantifying the proportion of shared detected loci relative to the union. Color intensity reflects the degree of overlap (from low to high). Cells labeled “NA” indicate comparisons where no loci were detected by one or both configurations.

These results indicate that under a diffuse tiny-effect architecture, widely used GWAS methods can generate apparently localized association peaks despite broadly distributed underlying signals. This pattern is most evident for BOLT-LMM and REGENIE_LOCO, which detected many more significant SNPs and loci than other approaches, but it is not unique to them. The limited overlap across methods further suggests that such peaks do not necessarily mark discrete local genetic effects. Rather, in this setting they may represent method-dependent summaries of diffuse polygenic signals shaped by livestock LD.

### Long-range LD induces signal spillover and inflation

The definition of null regions was guided by the empirical decay of linkage disequilibrium (Figure 1). Because LD remained appreciable over megabases, extended null windows were used to evaluate the propagation of association signals beyond effect regions.

Consistent with this expectation, association signal extended beyond effect windows and persisted in null regions, including central-null regions (Figure 6). Spillover-locus rates in central-null regions (Figure 6A) differed markedly across methods. At the genome-wide significance threshold (*P* ≤ 5 × 10⁻⁸), the proportion of central-null 500 kb windows containing at least one significant SNP was 30.7% for BOLT-LMM, 30.3% for REGENIE_LOCO, 2.9% for FarmCPU (NYC), 1.9% for FarmCPU (Kansas), 1.5% for fastGWA, 1.1% for BLINK (NYC), 1.1% for REGENIE_woLOCO, 1.0% for BLINK (Kansas), and 0.0% for SLEMM. Although SLEMM detected genome-wide significant loci within simulated effect windows (Figure 3), no central-null 500 kb window reached genome-wide significance in any replicate (Figure 6A). Thus, BOLT-LMM and REGENIE_LOCO exhibited extensive long-range propagation of LD-mediated association signals into effect-free regions. These results demonstrate that methods differ considerably in their susceptibility to long-range LD-driven false localization despite analyzing the same underlying polygenic architecture. Consistent patterns were observed for genomic inflation (Figure 6B). BOLT-LMM and REGENIE_LOCO exhibited the greatest inflation, with *λ_GC_* of 6.65–9.44 and 6.68–10.19, respectively, across the 10 simulated phenotypes. FarmCPU (2.92–3.61), fastGWA (2.37–2.61), REGENIE_woLOCO (1.82–2.72), and BLINK (1.55–2.01) showed intermediate to moderate inflation. In contrast, SLEMM remained the most conservative, with *λ_GC_* of 0.56–0.62, below the null expectation of 1, indicating deflated (conservative) statistics in these effect-free regions. Because these regions contain no causal variants, the persistence of signal cannot be a local reflection of nearby causal effects; rather, for the LOCO methods it is the predictable consequence of long-range LD with same-chromosome effect windows (Figure 8), so their *λ_GC_* > 1 reflects genuine block-level signal that is misleading for localization.

**Figure 6.**
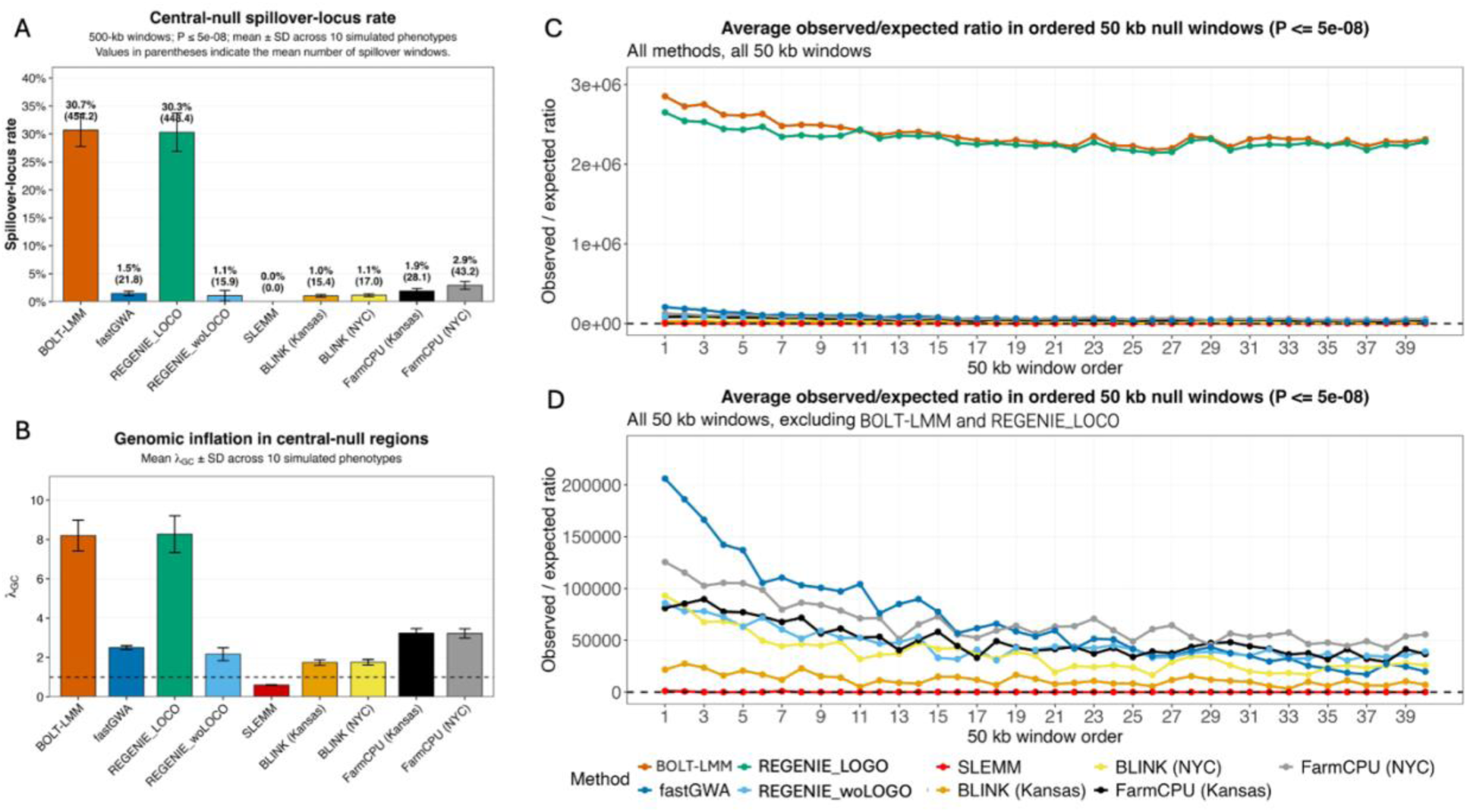
Long-range LD–induced spillover and spatial signal decay. (A) Spillover-locus rate in central-null regions. Each central 2 Mb null region was partitioned into consecutive 500 kb windows. A window was counted as a spillover locus if it contained at least one SNP reaching genome-wide significance (*P* ≤ 5 × 10⁻⁸). The spillover-locus rate was calculated as the fraction of central-null 500 kb windows containing a genome-wide significant association. Bars show the mean spillover-locus rate across 10 simulated phenotypes, error bars indicate ± SD. Labels above the bars indicate the mean spillover-locus rate (%) and the corresponding mean number of spillover loci (500 kb windows) per genome. (B) Genomic inflation factors (*λ_GC_*) estimated within central-null regions (2 Mb windows). Because these regions contain no causal variants but remain in long-range LD with same-chromosome effect windows, *λ_GC_* > 1 for the LOCO methods reflects genuine LD-aggregated block-level signal (Figure 8), whereas *λ_GC_* < 1 for SLEMM reflects absorption of that block-level signal into the full-GRM polygenic background. (C) Observed-to-expected ratios of significant SNPs across ordered 50 kb null windows for all methods, calculated as the number of observed significant SNPs divided by the expected number under the nominal significance threshold (the number of SNPs tested within each window multiplied by α = 5 × 10⁻⁸). Ratios greater than 1 indicate enrichment of significant associations relative to the null expectation and show strong and persistent inflation for methods with extensive LD propagation. (D) Same analysis excluding BOLT-LMM and REGENIE_LOCO to highlight differences among remaining methods. Together, these panels illustrate method-specific differences in the magnitude and spatial extent of LD-driven signal propagation.

Spatial analysis of ordered null windows further supported this interpretation (Figure 6C–D). Although signal generally decayed with distance from effect boundaries, BOLT-LMM and REGENIE_LOCO remained inflated across null windows, indicating persistent long-range LD propagation. The remaining methods showed clearer attenuation with distance, but still differed substantially in the extent to which signal persisted in null space, whereas SLEMM remained stable throughout.

Collectively, these results show that under a polygenic architecture, GWAS signal is not confined to effect windows but can propagate into distal null regions through long-range LD. The extent of this propagation depends strongly on method specification, indicating that different GWAS approaches handle long-range LD very differently.

### Peak loci are unstable across phenotypes

We quantified the overlap of significant 500 kb loci across phenotype replicates using the Jaccard index (Figure 7). Despite shared causal locations and generating distributions, most methods showed very low overlap, indicating that detected peak locations were highly unstable. BLINK and FarmCPU exhibited consistently low overlap (generally ∼0.04–0.08), while fastGWA showed only modestly higher consistency (∼0.06–0.13). In contrast, BOLT-LMM and REGENIE_LOCO achieved moderate but still limited overlap (∼0.19–0.35), suggesting some reproducibility but far from stable localization. REGENIE_woLOCO showed substantially reduced consistency, with overlaps largely near zero (∼0.00–0.08). SLEMM exhibited essentially no overlap because it detected almost no loci. Overall, these results demonstrate that the genomic positions of significant loci vary substantially across replicates, indicating that detected peaks do not represent stable, well-localized features of the underlying polygenic signal.

**Figure 7.**
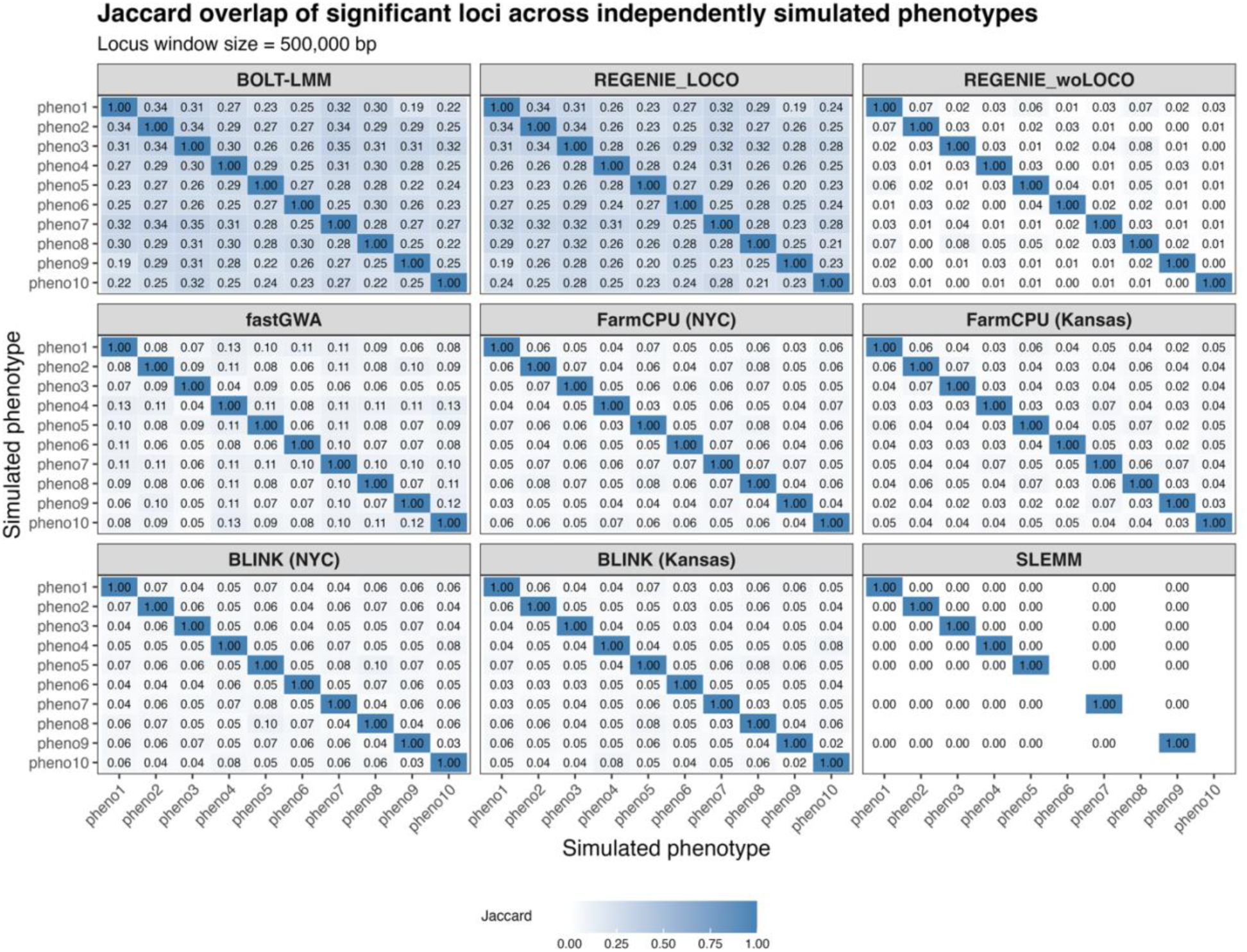
Instability of GWAS peak loci across phenotype replicates. Jaccard overlap of significant 500 kb loci between phenotype replicates for each method configuration. Each panel shows pairwise overlap of detected loci across 10 simulated phenotypes sharing the same underlying polygenic architecture.

### LOCO test statistics quantitatively match the LD-aggregated polygenic prediction

We compared the observed GWAS test statistics with the a priori prediction from Equation 13 of Jiang^54^. The 135,358 SNPs were grouped into 20 equal-count bins according to the predicted non-centrality parameter (NCP), and the observed mean χ² across 10 phenotype replicates was calculated for each bin. For both BOLT-LMM and REGENIE_LOCO, the observed mean χ² closely followed the predicted expectation across the full range of NCP values (Figure 8), indicating that the elevated test statistics observed for the LOCO methods are quantitatively explained by LD-aggregated polygenic signal.

**Figure 8.**
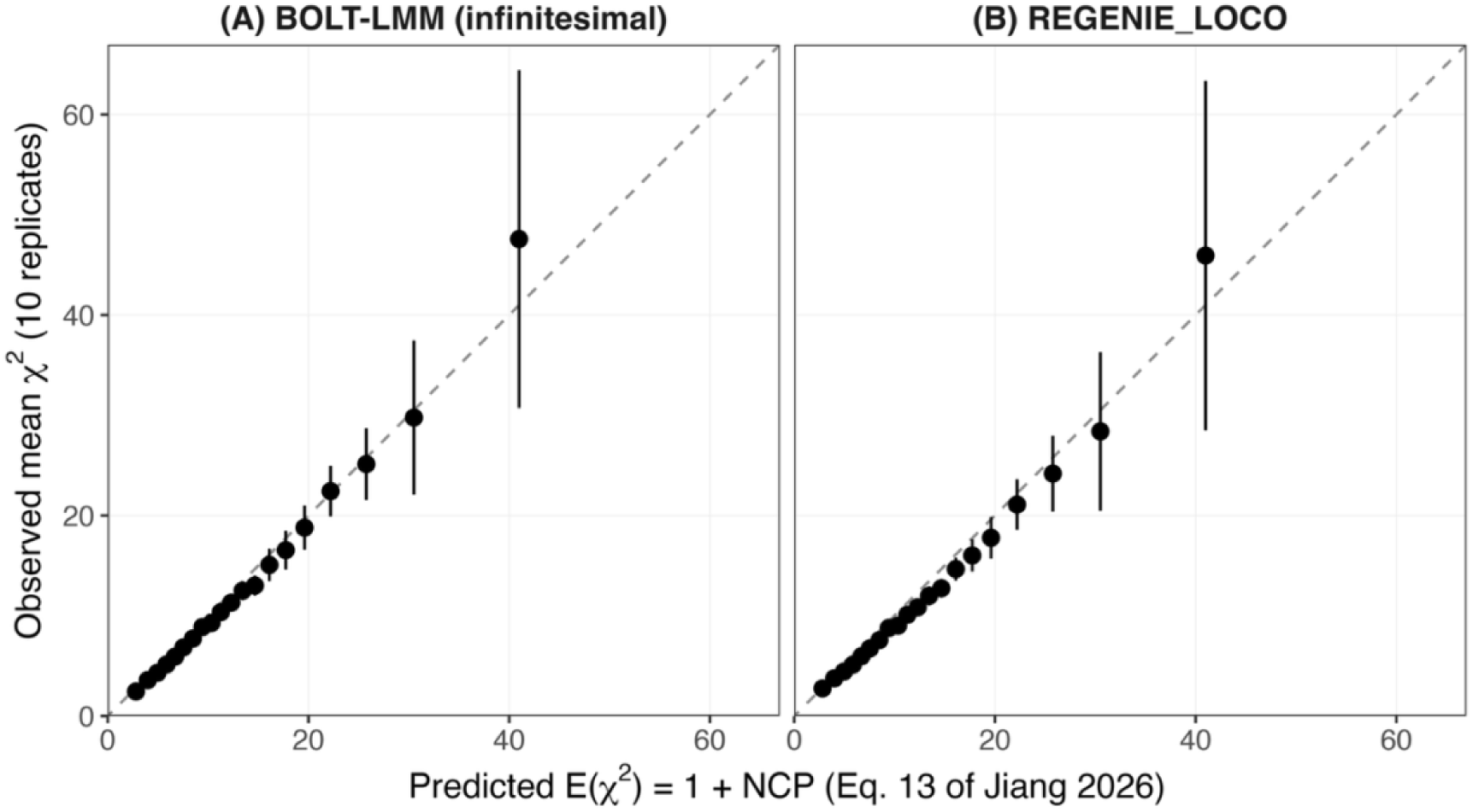
Comparison of the theoretical prediction from Equation 13 of Jiang ^54^ with the observed GWAS test statistics. Panel (A) shows BOLT-LMM using the infinitesimal-model statistic, and panel (B) shows REGENIE_LOCO. The 135,358 SNPs were divided into 20 equal-count bins according to the theoretical non-centrality parameter (NCP) predicted by Equation 13, using only the theoretical prediction and without reference to any GWAS statistic. For each bin, the x-coordinate is the mean predicted test statistic, E(χ²) = 1 + NCP, and the y-coordinate is the observed mean χ² across all SNPs in the bin and 10 independently simulated phenotype replicates. Error bars represent 95% confidence intervals calculated from variation among the 10 replicate-specific bin means as 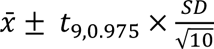 requiring no assumption of independence among SNPs, genomic blocks, or chromosomes. The inputs c_LOCO, the causal LD score (L_j), and the per-SNP effect variance (*h*²/*m*) were obtained from the genotypes and the known simulated architecture, with no parameters fitted to the GWAS results. The dashed line is the identity line (y = x), representing perfect calibration rather than a fitted regression.

Taken together, these results show that GWAS methods differ markedly in how diffuse polygenic signals are transformed into variant-level associations under realistic LD structure. BOLT-LMM and REGENIE_LOCO produced the strongest signal amplification, yielding many significant associations but also substantial spread into null regions. At the other extreme, SLEMM showed the strongest spillover control, with deflated null-region statistics, whereas fastGWA, REGENIE_woLOCO, BLINK, and FarmCPU showed intermediate behavior.

Importantly, apparent detection in this benchmark should not be interpreted as accurate localization. Under this diffuse polygenic architecture, significant associations frequently extended beyond effect windows and appeared in null and central-null regions, indicating that prominent GWAS peaks often reflect method-dependent LD aggregation rather than recovery of discrete causal loci. Overall, these findings show that GWAS methods can differ not only in power, but also in the extent to which they convert distributed polygenic background into apparently localized signals. This underscores the importance of evaluating GWAS methods in spatially explicit frameworks that account for LD structure and highly polygenic architectures, particularly in livestock populations with extensive long-range LD.

## Discussion

The central finding of this benchmark is not that GWAS fails under polygenic architecture, but that apparent localization can be deeply misleading in livestock-like genomes with extended linkage disequilibrium (LD). Because the simulated genetic signal was distributed across many small-effect variants rather than concentrated in discrete major loci, statistically significant associations cannot be interpreted as evidence of a nearby causal variant of meaningful effect. This follows directly from the statistical conditions for detection in GWAS. The single-marker test statistic satisfies approximately *χ*^2^ ≈ 1 + *Nq_j_*, where *N* is sample size and *q_j_* is the proportion of variance explained (PVE) by SNP *j*. Genome-wide significance *P* = 5 × 10⁻^8^ corresponds to *χ*^2^ ≈ 30; at *N* = 20,000, a SNP must explain approximately *q_j_* ≳ 5 × 10⁻^3^ of phenotypic variance to be individually detectable at 50% power. In our simulation, 10,000 SNPs collectively explain a total heritability of 0.5, placing even the max per-SNP PVE substantially below this threshold; essentially none is expected to reach genome-wide significance on its own. The many significant associations reported by LOCO, fastGWA, and the multi-locus methods cannot be reconciled with such an individual-effect expectation and must instead arise from polygenic confounding through LD aggregation. The critical question is therefore not why individual SNPs fail to reach significance, but why many nevertheless do. The answer lies in LD-mediated aggregation. Under the marginal model, if the true architecture is *y* =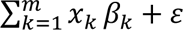, the estimated effect at marker *j* is approximately 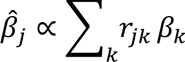, where r*j_k_* denotes LD between marker *j* and causal variant *k*. Even when each *β_k_* is individually negligible, the cumulative contribution of correlated variants within an LD block can push the marginal statistic at a tagging SNP well above the significance threshold. In a livestock genome with long-range LD, these blocks are broad, meaning that a single peak may summarize the collective signal of dozens to hundreds of weakly correlated variants spread across a genomic segment far wider than any fine-mapping window. Prominent GWAS peaks therefore do not mark nearby moderate-effect loci; they mark the centers of LD-mediated aggregation zones, and their apparent sharpness is a property of correlation structure rather than of underlying effect size architecture. The empirical results confirm this mechanism: spillover-locus rates in central-null regions, genomic inflation in central-null regions, and the spatial extent of observed-to-expected excess across ordered null windows all reflect the same underlying process of diffuse signal being concentrated and propagated through LD, with the magnitude of each phenomenon varying markedly across methods. This mechanistic interpretation is further supported by the close agreement between the theoretical prediction from Equation 13 of Jiang^54^ and the observed LOCO test statistics, indicating that the magnitude of the apparent localized associations is quantitatively consistent with LD-aggregated polygenic signal.

A second major result is that this behavior is strongly method-dependent, and the differences are traceable to concrete algorithmic choices rather than to power or sample size alone. BOLT-LMM and REGENIE_LOCO produced the largest numbers of significant SNPs and loci, the broadest signal profiles, and the greatest propagation into null space, making them the most prone in this benchmark to converting diffuse polygenic background into apparently localized associations, primarily because both employ a leave-one-chromosome-out scheme in which the background model is built from all chromosomes except the one being tested. Although this design prevents proximal contamination (the absorption of genuine local signal into the background term), it creates a systematic gap in polygenic control under diffuse architecture: the chromosome-specific polygenic variance omitted from the LOCO predictor is approximately proportional to the number of causal variants assigned to each chromosome rather than being uniform across chromosomes. In our simulation, the expected chromosome-specific PVE ranged from 0.0138 (chromosome 12) to 0.0497 (chromosome 1), inflating test statistics across the excluded chromosome in proportion to each SNP’s LD score (its local and long-range LD with effect-window variants) and producing signal that is strongest near effect windows but persists into null and central-null regions far from any effect region. Because the infinitesimal BOLT-LMM statistic (P_BOLT_LMM_INF) was analyzed, the similarity between BOLT-LMM and REGENIE_LOCO is attributable primarily to their shared LOCO strategy rather than to BOLT-LMM’s optional non-infinitesimal mixture prior. fastGWA, BLINK, FarmCPU, and REGENIE_woLOCO showed intermediate behavior for related but distinct reasons: fastGWA retains only sparse GRM elements above a kinship threshold, leaving residual LD-aggregated signal that a dense full GRM would absorb; BLINK and FarmCPU replace the continuous polygenic random effect with iteratively selected marker cofactors that cannot adequately capture background variance under a diffuse, highly polygenic architecture; and REGENIE_woLOCO, by including the tested chromosome in its ridge regression predictor, partially absorbs the focal chromosome’s polygenic contribution (which under diffuse architecture improves calibration relative to the LOCO version, though residual inflation persists because the predictor remains an approximation rather than a full dense GRM). SLEMM was substantially more conservative, producing no genome-wide significant loci in central-null regions, exhibiting deflated central-null test statistics (*λ_GC_* = 0.56-0.62), and remaining stable across ordered null windows, because its full dense GRM without LOCO exclusion correctly absorbs the aggregate of tiny effects as background variance, eliminates the per-chromosome control gap, and avoids the cumulative upward biases introduced by stochastic or block-based approximations used in other methods. The contrast between REGENIE with and without LOCO is especially informative: it demonstrates that the phenomenon is not an inevitable consequence of the data structure alone, but depends materially on how a method partitions background polygenic signal during testing, a difference of a single algorithmic switch that shifts a method from intermediate to maximally inflated behavior.

A subtler reading of these method-dependent differences is that they reflect different inferential targets rather than different degrees of error. Under the SNP-level inference that motivates most downstream GWAS analyses (candidate-gene prioritization, fine-mapping, and colocalization), the spillover signals reported in null and central-null regions by LOCO and sparse-GRM methods are misleading peaks: they do not localize to any particular nearby variant and cannot be honestly followed up at single-marker resolution. Under a block-level reading, however, those same signals faithfully reflect long-range LD with on-chromosome effect-window variants that LOCO cannot absorb because it excludes the focal chromosome from the polygenic background. Both readings can be simultaneously true: LOCO is detecting genuine LD-aggregated block-level signal, while full-GRM is restricting attention to SNP-level excess above the polygenic baseline ^54^. For livestock GWAS aimed at SNP-level interpretation, the full-GRM reading is the operationally relevant one, and SLEMM’s conservatism is therefore appropriate to that goal rather than a power deficit.

The heritability estimation results provide an additional mechanistic lens through which the intermediate behavior of fastGWA can be understood. The consistent underestimation of 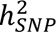 by fastGWA-REML (recovering approximately 29% of the simulated heritability despite a true value of 0.5) directly reflects the consequence of GRM sparsification for background polygenic control. When the GRM retains only elements above a kinship threshold, the realized genomic relationships among distantly related individuals, which collectively carry the bulk of the polygenic signal under a diffuse, highly polygenic architecture, are set to zero and excluded from the variance component model. The estimated genetic variance therefore captures only the component attributable to close relative pairs, leaving the remainder unmodeled. This has a direct downstream consequence for association testing: the polygenic background term fitted during GWAS is calibrated to an underestimated heritability, meaning that a substantial fraction of the true polygenic variance is not absorbed into the random effect and instead remains in the residual, where it can inflate single-marker test statistics through LD aggregation. The intermediate spillover-locus rates and null-window inflation observed for fastGWA are thus a predictable consequence of this incomplete background modeling, and the degree of inflation should be expected to scale with the gap between the true heritability and the heritability captured by the sparse GRM. More broadly, this result illustrates that the choice of GRM approximation has consequences not only for computational efficiency but for the statistical validity of the association test itself, a consideration that is particularly consequential in livestock populations where highly polygenic architectures and extended LD jointly amplify the effects of incomplete background control.

The instability of detected loci across phenotype replicates further strengthens this interpretation. Although all replicates shared the same underlying polygenic architecture and population LD structure, overlap of significant 500 kb loci across replicates was generally low, particularly for BLINK, FarmCPU, fastGWA, REGENIE_woLOCO, and SLEMM. Even BOLT-LMM and REGENIE_LOCO, which showed the greatest consistency, achieved only moderate overlap despite producing the largest absolute numbers of significant loci. This pattern is diagnostic: if detected peaks reflected stable, well-localized effects driven by the underlying genetic architecture, reproducibility across replicates sharing that architecture should be substantially higher. The observed low and method-dependent overlap instead indicates that many significant loci are contingent on the particular realization of the small effects through LD and method-specific background modeling, rather than being robust features of the architecture itself. Figure 7 therefore measures the stability of detected loci across independent realizations of this diffuse architecture (fixed causal locations, with effects and residuals redrawn). The implication extends beyond this benchmark: cross-study concordance of peaks produced by the same method in livestock populations should not be taken as strong evidence of true localization, as shared LD structure and shared method choice can produce concordant spurious peaks independently of shared underlying biology.

This interpretation is compatible with broader polygenic and omnigenic conceptualizations of complex trait architecture, in which weak effects distributed widely across the genome become statistically concentrated through LD correlation structure, though the benchmark does not test or prove the omnigenic model itself. The livestock context deserves specific note: smaller effective population sizes, stronger artificial selection, and more extended long-range LD relative to outbred human populations may make LD-mediated aggregation systematically more severe in livestock GWAS than in human GWAS conducted under comparable sample sizes. Peaks that would reflect genuine moderate-effect loci in a human GWAS context may, in a livestock genome, be predominantly LD-aggregation artifacts of the kind demonstrated here, and this distinction has practical consequences for how the existing livestock GWAS literature should be read, particularly for traits with highly polygenic architectures where detected peaks from liberal methods may warrant reinterpretation rather than direct follow-up.

Several limitations of the present benchmark should be acknowledged. The simulation used a single idealized infinitesimal architecture in which causal SNP effects were drawn from a single normal distribution, each explaining on average 0.005% (0.5/10,000) of phenotypic variance; real traits likely have architectures intermediate between infinitesimal and oligogenic, with moderate-effect variants embedded in a polygenic background, and the relative performance ordering of methods could shift under such mixed architectures. The benchmark also used a single livestock population LD structure, and the severity of LD-mediated aggregation will depend on the LD decay characteristics of the specific breed or species analyzed. The sample size of *N* = 20,000 is realistic for some well-resourced livestock populations but exceeds what is available for many breeds, and at smaller sample sizes the absolute number of spurious detections would decrease, though the relative ordering of methods and the underlying mechanism would remain the same. Finally, SLEMM’s conservatism is a virtue under a diffuse, highly polygenic architecture but carries a genuine cost under architectures with moderate-effect loci mixed into a polygenic background, where the full-GRM approach can suppress genuine signals through proximal contamination; appropriate method choice therefore depends on prior expectations about the trait architecture under study.

The primary practical implication of this benchmark is a caution about interpretation that goes beyond simply choosing a more conservative method. In standard GWAS practice, a significant peak is typically treated as evidence for a nearby variant of meaningful effect, a starting point for fine-mapping, candidate gene identification, colocalization, or functional follow-up. However, the results here demonstrate that significant peaks can arise from at least three qualitatively distinct sources: a genuine nearby variant with a large or moderate effect, the LD-mediated accumulation of many tiny polygenic effects across a broader genomic segment, or outright false positives driven by incomplete polygenic background control. These three sources are not equivalent in their value for downstream analysis. A peak anchored by a large-effect variant retains meaningful localization value (the associated genomic region genuinely points toward a functional mechanism, and fine-mapping, colocalization, and candidate gene analyses are well-motivated). A peak arising from the accumulation of small polygenic effects across a broad LD block, by contrast, may not resolve to a single nearby causal variant. Instead, it summarizes diffuse signal whose spatial origin spans the LD segment rather than any particular locus within it, limiting the precision with which specific causal variants or genes can be prioritized. As recently demonstrated by Jiang^54^, long-range LD fundamentally limits the resolution of GWAS fine-mapping, making it difficult to accurately identify the true causal variant. Therefore, these peaks should not be interpreted as direct evidence for a specific causal variant. Rather, they should be regarded as regional association signals that require additional evidence, such as functional annotation, experimental validation, or complementary genomic analyses, before a more refined search for the causal variant is undertaken. A false-positive peak offers nothing beyond noise. The practical difficulty is that these three types of peaks can appear visually indistinguishable in a Manhattan plot (all three can produce sharp, prominent associations), yet they demand fundamentally different downstream responses. Distinguishing peaks that harbor genuine large-effect variants from those that reflect polygenic accumulation or false positives is therefore not a refinement of GWAS interpretation but a prerequisite for it, particularly in livestock populations where extended LD makes polygenic accumulation peaks both more common and more visually convincing. Toward this end, several complementary diagnostics are available to practitioners. Comparing results with and without LOCO, or across full-GRM and sparse-GRM implementations, provides a first filter: substantial disagreement in the number and spatial extent of significant loci between these versions is more consistent with LD-mediated inflation than with stable large-effect detection. Checking locus stability across phenotype replicates or bootstrap samples provides a second filter, as peaks driven by genuine large-effect variants should reproduce consistently across replicates sharing the same architecture, whereas peaks arising from polygenic accumulation or false positives will be more replicate-dependent. Examining the per-SNP effect size distribution within a significant window (whether the peak is anchored by one or a few SNPs with substantially larger marginal effects than their neighbors, or instead shows a flat, diffuse elevation across the window) offers a third diagnostic, as genuine large-effect loci tend to produce a discernible lead SNP while polygenic accumulation peaks tend to be broad and undifferentiated. When the primary goal is biologically meaningful localization rather than simply detecting variant-level associations, our benchmark suggests that more conservative full-GRM approaches such as SLEMM provide a more reliable foundation in livestock-like settings by suppressing polygenic accumulation peaks and retaining primarily those associations with sufficient per-locus signal to survive complete background control. Peaks identified by more liberal methods should be accompanied by explicit acknowledgment that they may represent LD-aggregated summaries of diffuse polygenic signal, and prioritization of peaks for downstream follow-up should incorporate stability, effect size concentration, and method-concordance criteria rather than relying on significance alone.

## Supporting information

Supplementary_Figures_S1-S10_Manhattan_Plots

## Author contributions

J.J. conceived and designed the study and contributed the theoretical non-centrality-parameter analysis. X.W. and J.W. conducted the benchmarking analyses. F.T., Y.H., and C.M. contributed to data preprocessing. F.T., W.H., and C.M. contributed to interpretation of the results. X.W. wrote the initial draft, and J.J. revised the manuscript. All authors reviewed and approved the final manuscript.

## Data Availability

The individual-level pig genotype data used in this study are proprietary to Smithfield Premium Genetics and are not publicly available. Access to these data may be available from Smithfield Premium Genetics upon reasonable request and subject to a data-sharing agreement and any applicable institutional approvals. The analysis code used for phenotype simulation, GWAS benchmarking, and the theoretical non-centrality parameter calculations, together with derived summary statistics, is publicly available at https://github.com/xwang445/livestock-gwas-benchmark.

