## Supplementary_Figures_S1-S10_Manhattan_Plots for "Widely used GWAS methods can be poorly suited to SNP-level localization under diffuse polygenic architecture in livestock"

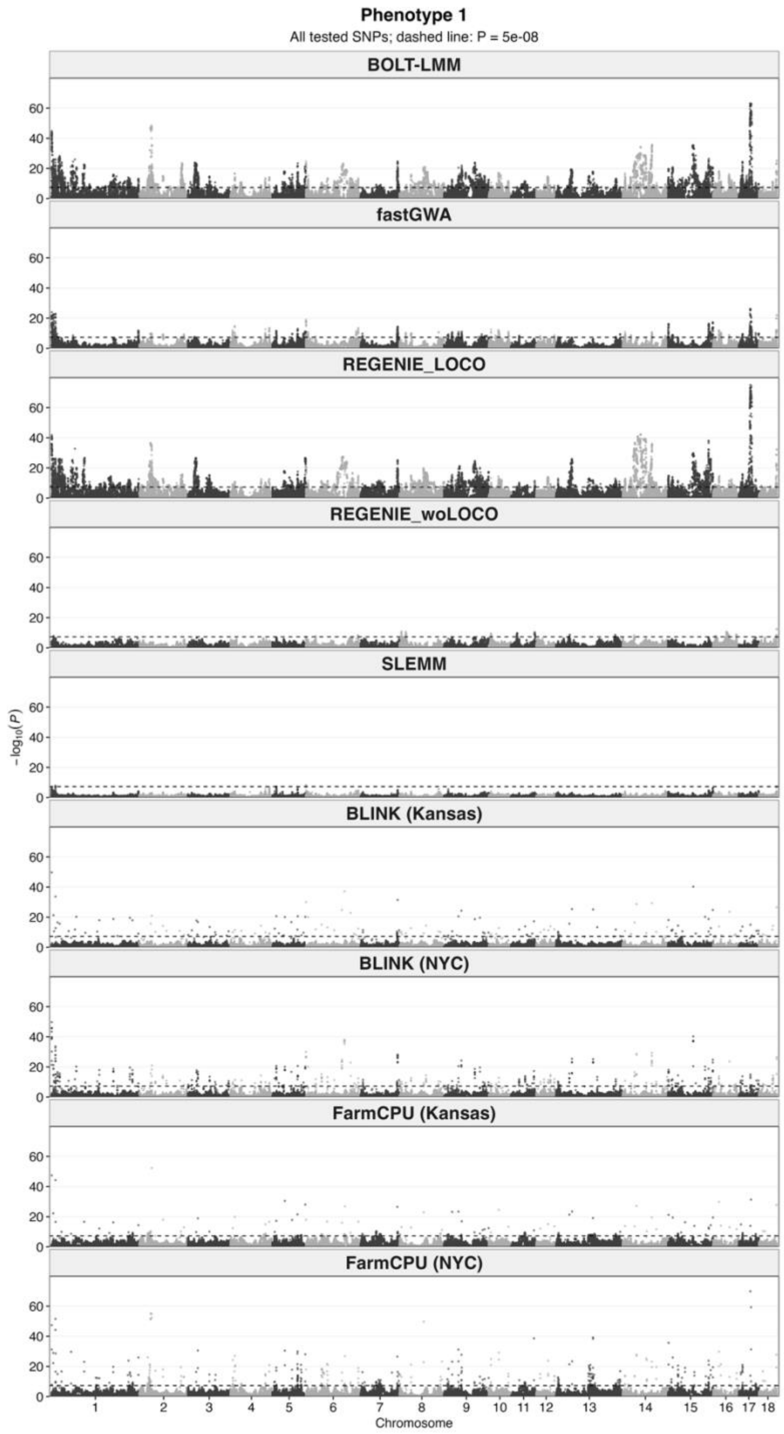


**Supplementary Figure S1.** Manhattan plots for **simulated phenotype 1** across all nine GWAS method configurations. Each panel represents one GWAS method configuration and displays all tested SNPs according to their genomic positions. Alternating shades distinguish adjacent chromosomes. The dashed horizontal line indicates the genome-wide significance threshold ($P = 5 \times 10^{-8}$). REGENIE_LOCO and REGENIE_woLOCO denote REGENIE with and without leave-one-chromosome-out, respectively.

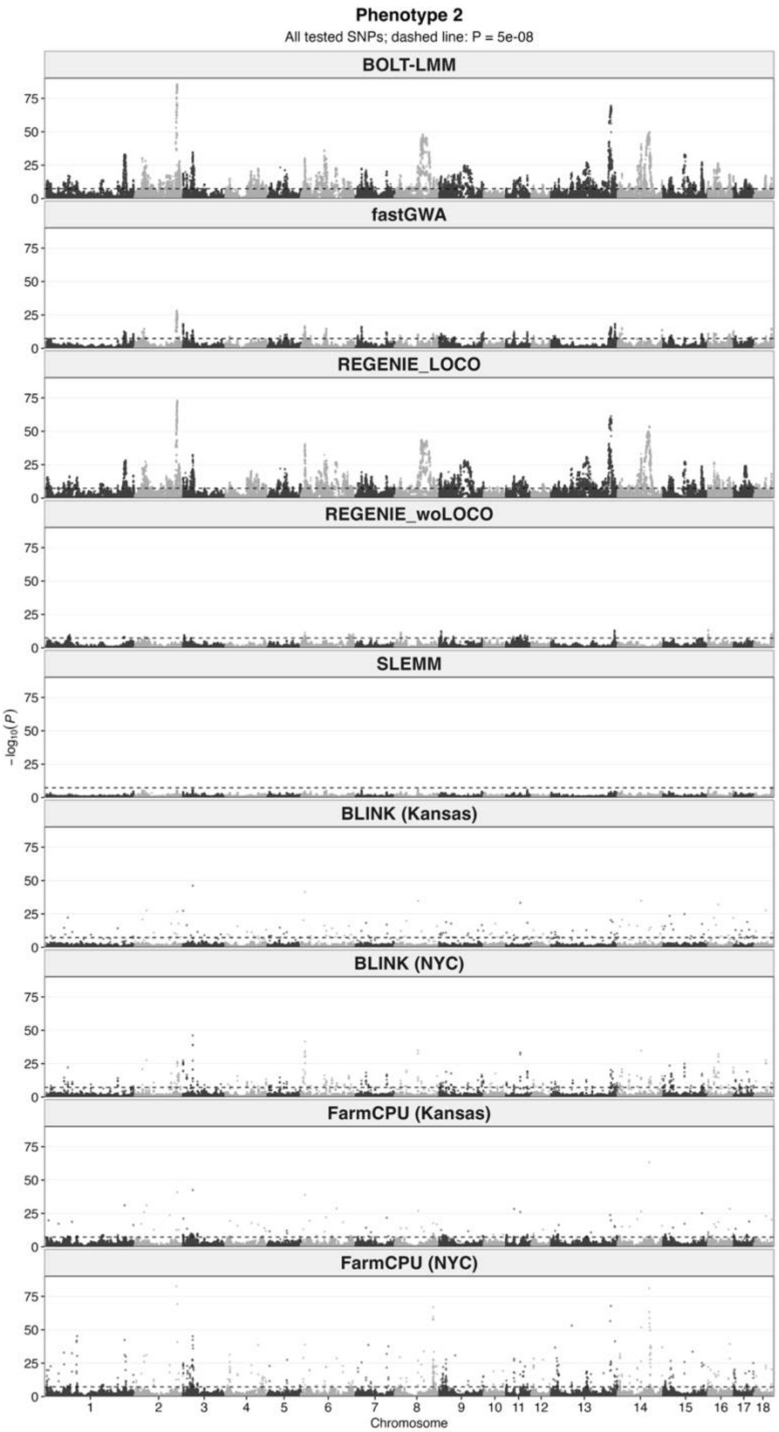


**Supplementary Figure S2.** Manhattan plots for simulated phenotype 2. See Supplementary Figure S1 for plotting details.

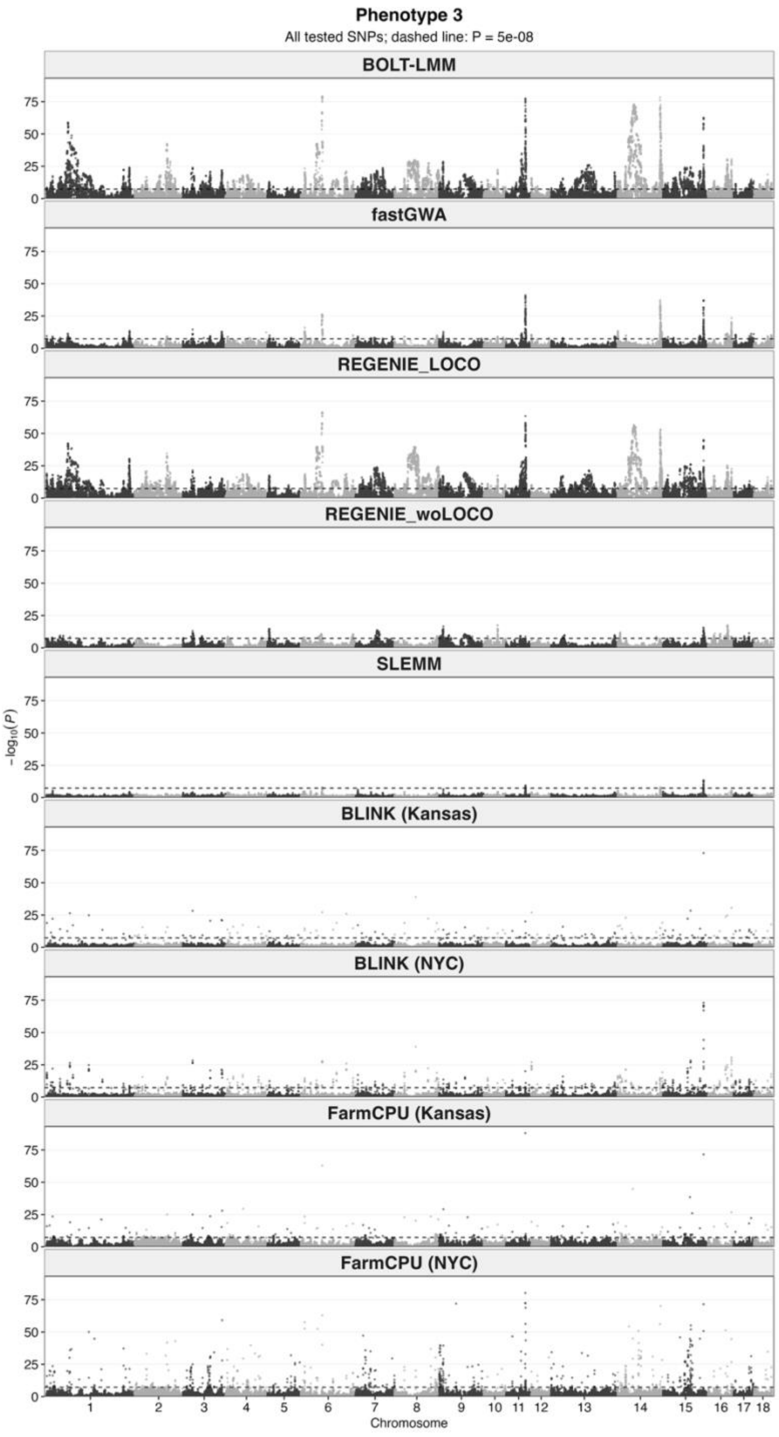


**Supplementary Figure S3.** Manhattan plots for simulated phenotype 3. See Supplementary Figure S1 for plotting details.

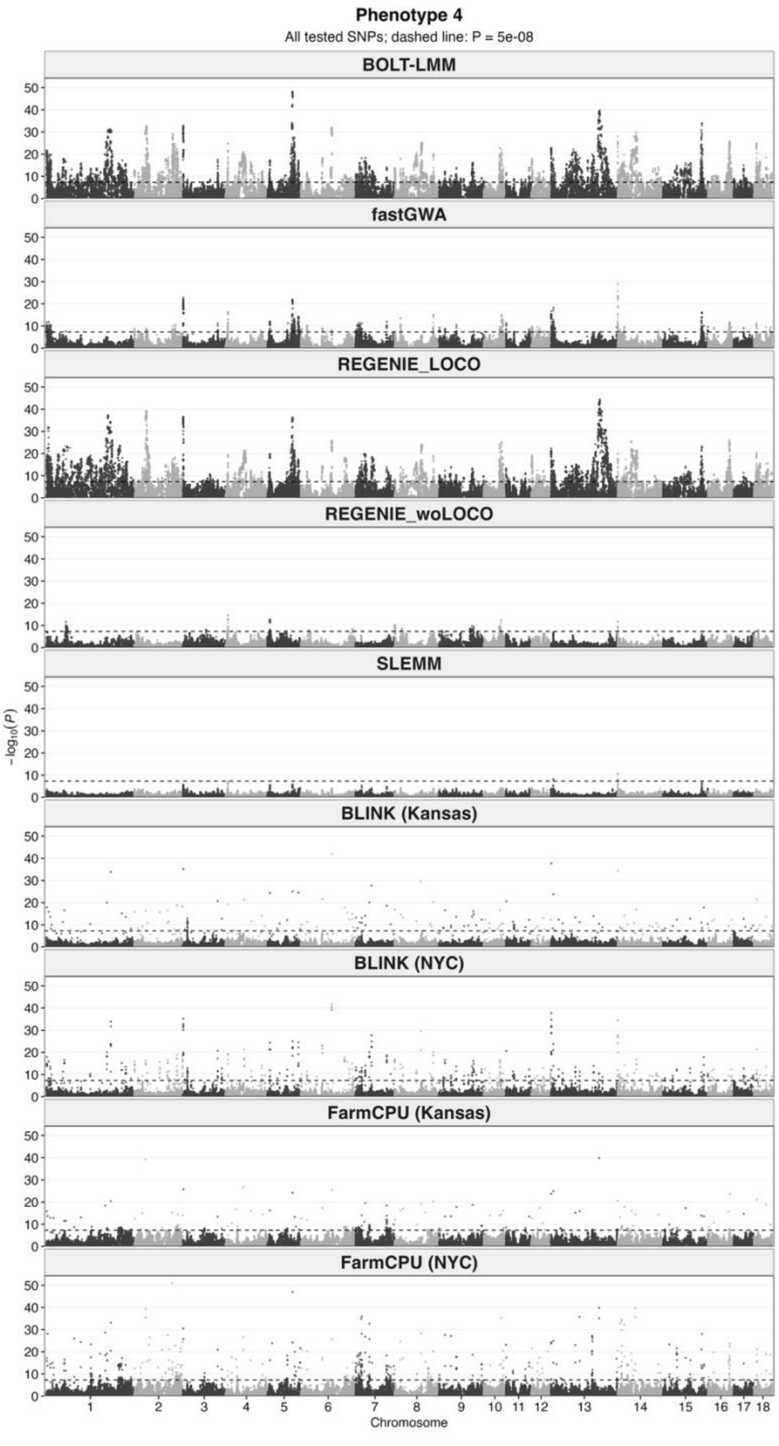


**Supplementary Figure S4.** Manhattan plots for simulated phenotype 4. See Supplementary Figure S1 for plotting details.

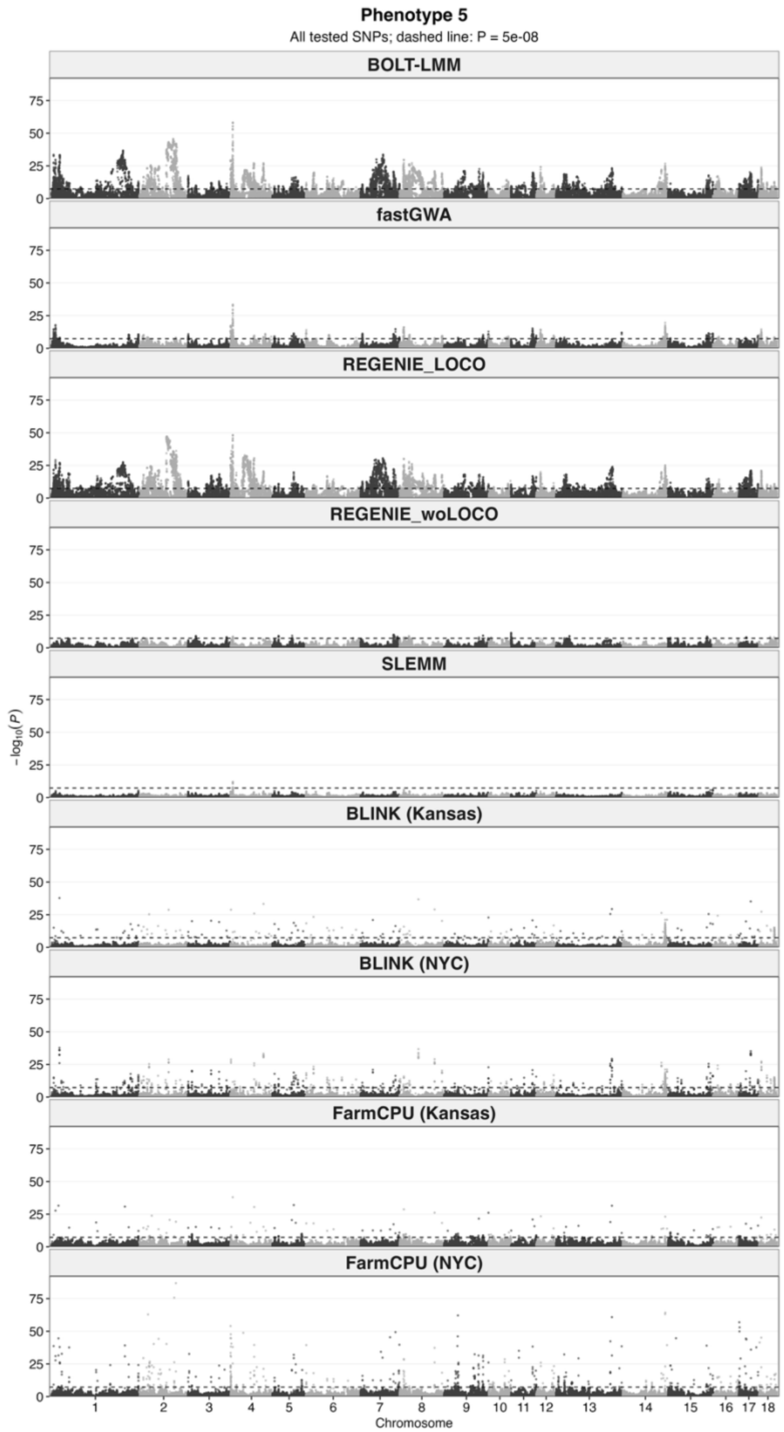


**Supplementary Figure S5.** Manhattan plots for simulated phenotype 5. See Supplementary Figure S1 for plotting details.

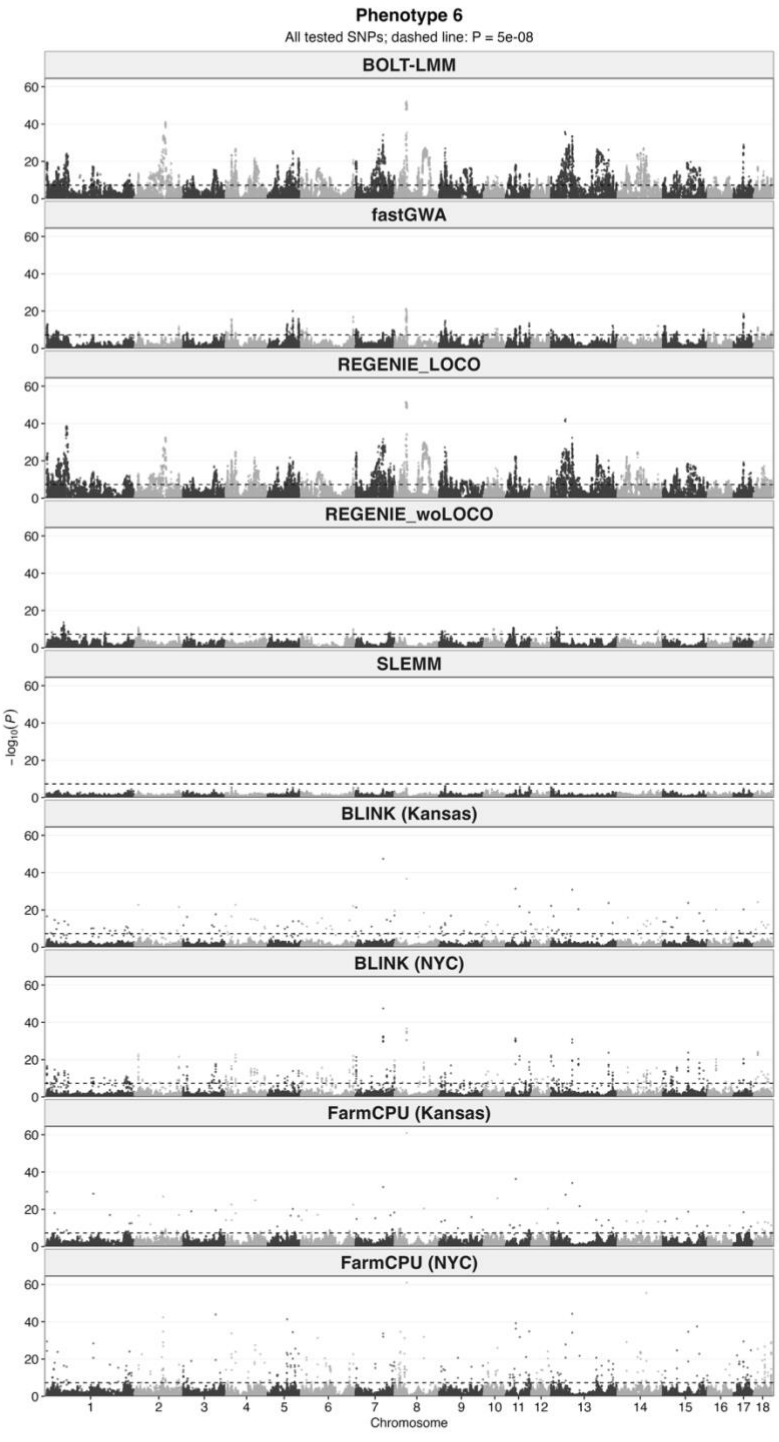


**Supplementary Figure S6.** Manhattan plots for simulated phenotype 6. See Supplementary Figure S1 for plotting details.

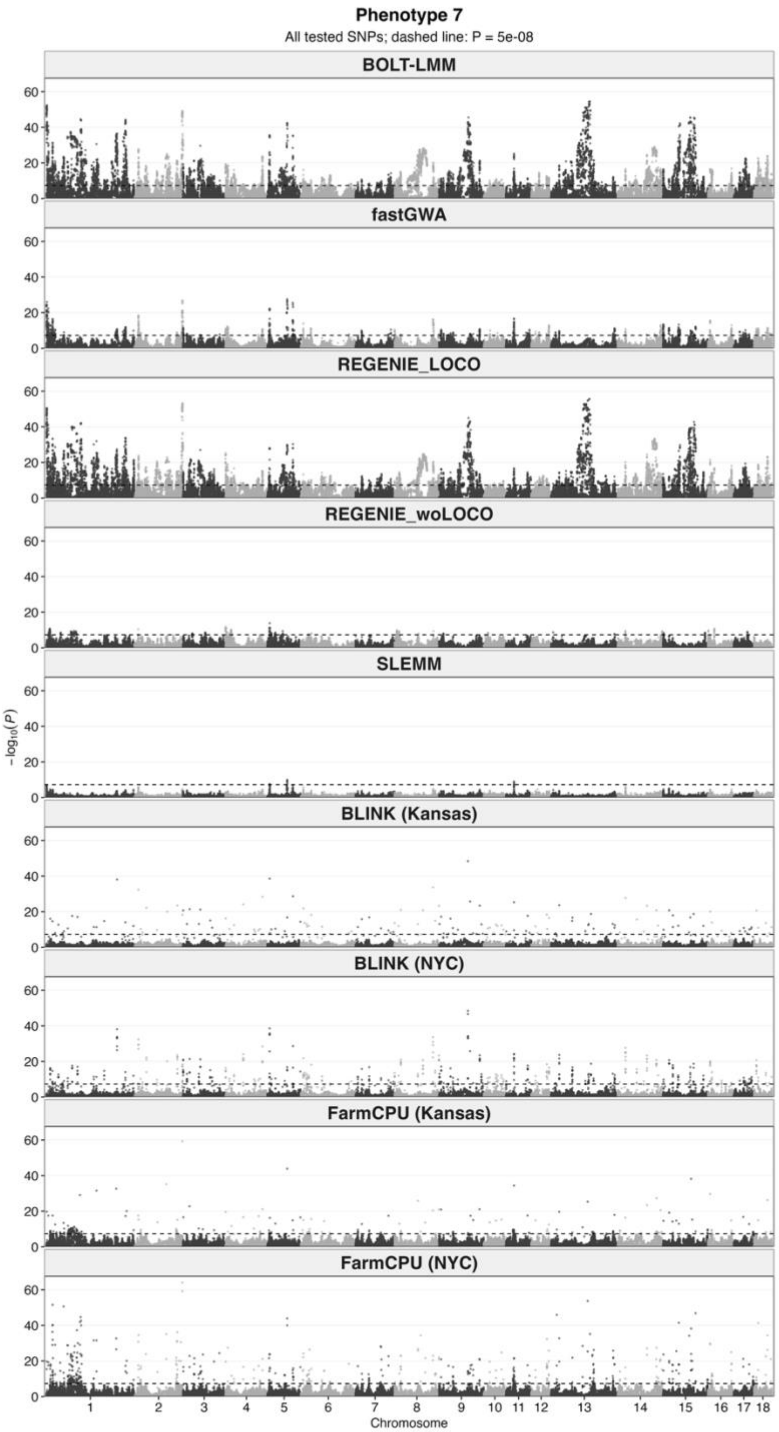


**Supplementary Figure S7.** Manhattan plots for simulated phenotype 7. See Supplementary Figure S1 for plotting details.

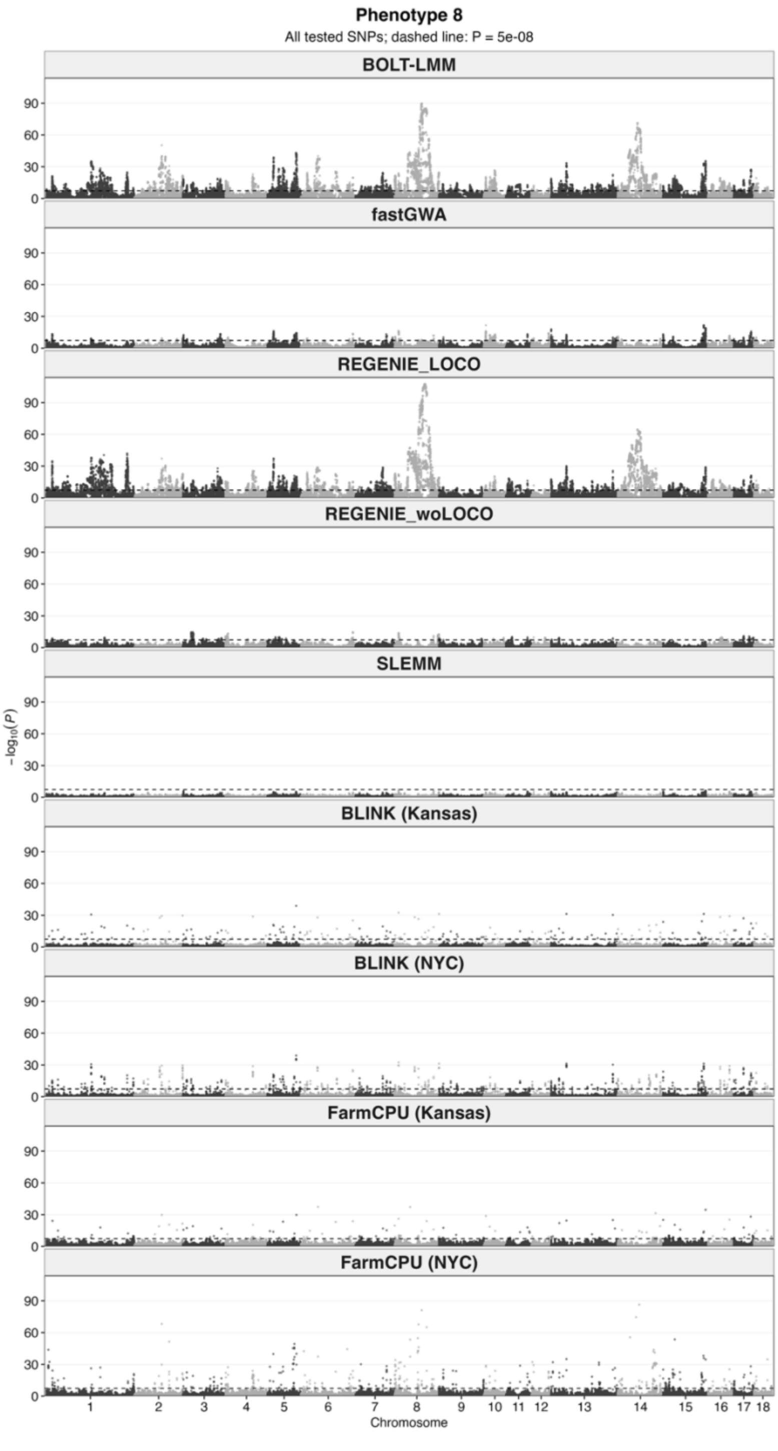


**Supplementary Figure S8.** Manhattan plots for simulated phenotype 8. See Supplementary Figure S1 for plotting details.

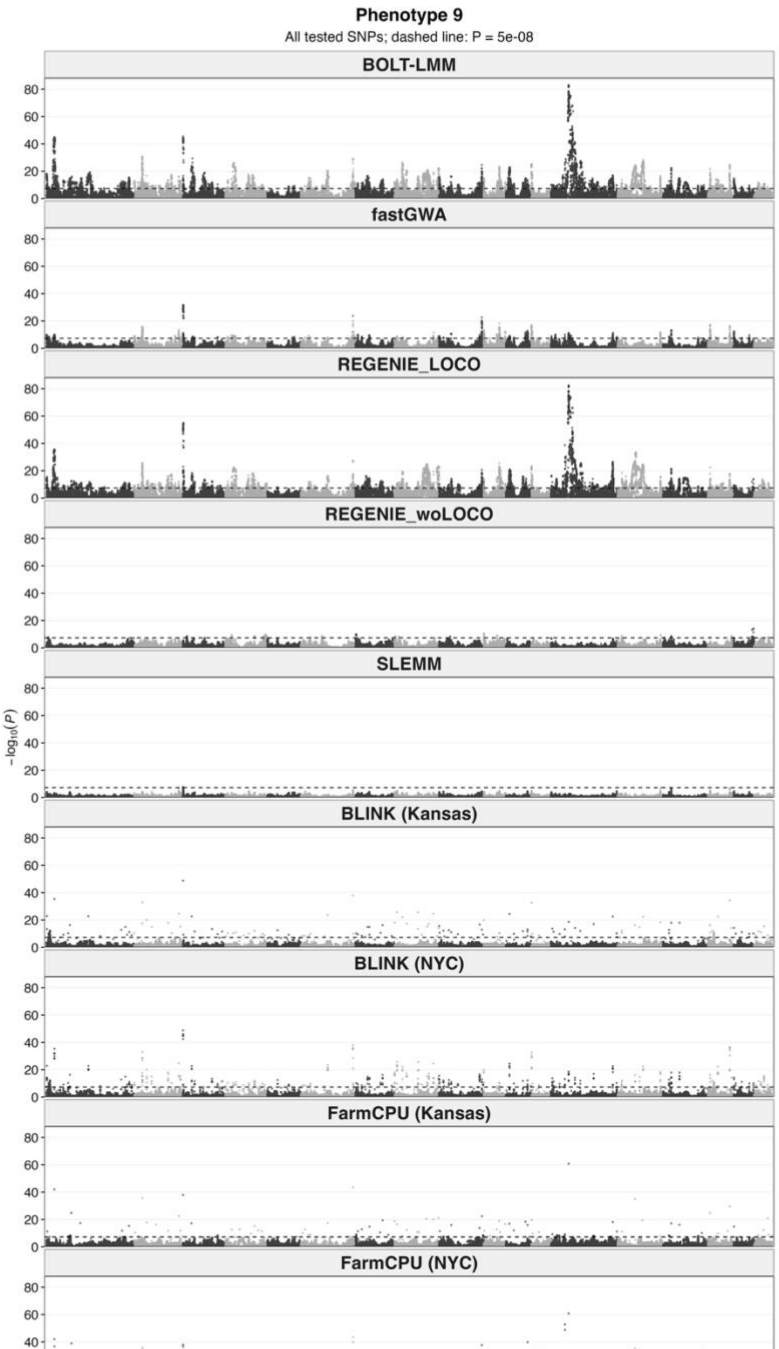


**Supplementary Figure S9.** Manhattan plots for simulated phenotype 9. See Supplementary Figure S1 for plotting details.

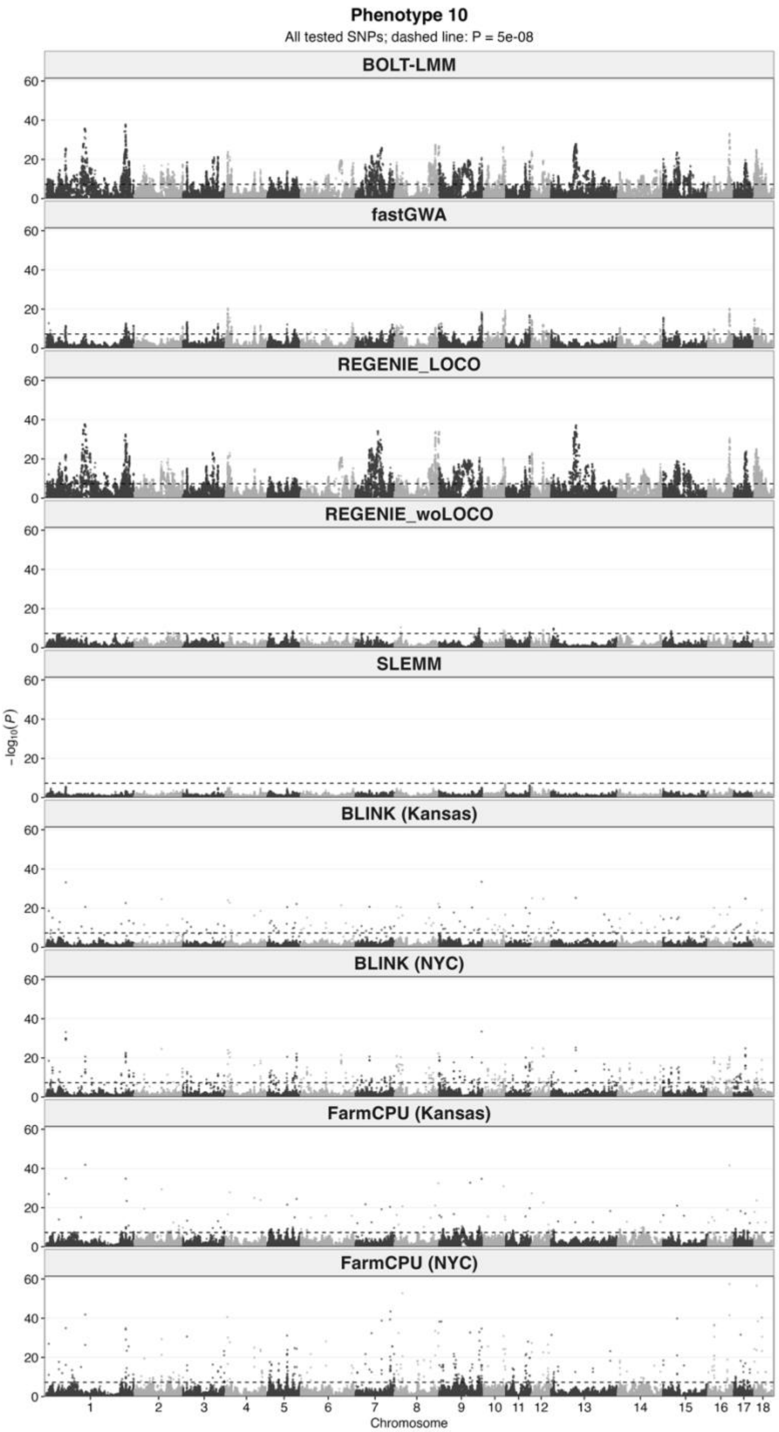


**Supplementary Figure S10.** Manhattan plots for simulated phenotype 10. See Supplementary Figure S1 for plotting details.
